# Nuclear effects play an important role in determining codon usage-dependent human gene expression

**DOI:** 10.1101/2024.10.20.619315

**Authors:** Renu Garg, Pancheng Xie, Jiabin Duan, Huan Liu, Yi Liu

## Abstract

Codon usage biases play a significant role in determining gene expression levels. Codon usage was thought to primarily influence translation-dependent processes. Here we show that codon usage optimality correlates genome-wide with nuclear mRNA and transcription levels in human cells, suggesting a nuclear role for codon usage. A genome-wide non-biased CRISPR-Cas9 screen was performed to identify factors involved in codon usage effects on gene expression. The identified factors include the CCR4-NOT complex subunit CNOT4 and many nuclear factors, including nuclear RNA exosome, PAXT complex components, and transcription factors. The CCR4-NOT complex has been shown to affect codon usage-dependent co-translational mRNA decay in yeast. Surprisingly, human CNOT4 was found to be highly enriched in nucleus and its influence on codon usage-dependent gene expression is largely caused by its impact on nuclear RNA levels due to transcriptional effect. On the other hand, nuclear exosome and PAXT complex, regulates nuclear mRNA stability through the RNA quality control pathway, leading to preferential cytoplasmic accumulation of mRNAs with optimal codon usage. Overall, our findings suggest that codon usage biases are due to selection on both translation-dependent and independent processes; different nuclear mechanisms play a key role in determining nucleotide composition-dependent gene expression levels in human cells.

## Introduction

Codon usage bias, the preference for certain synonymous codons for almost all amino acids, is a fundamental feature of eukaryotic and prokaryotic genomes. Although synonymous codon changes were previously thought to be silent, gene codon usage has been shown to play a major role in regulating gene expression and protein biogenesis in diverse organisms ^1–4^. In various organisms, genome-wide correlations between codon usage bias and protein levels have been observed ^5–7^. As expected from its role in encoding amino acids, the effects of codon usage on gene expression were thought to be mainly due to influences on translation-dependent processes. In support of this, codon usage has been shown to control translation elongation speed and subsequently affects co-translational protein folding ^8–14^. Due to the translational pausing at non-optimal codons during elongation, codon usage affects mRNA translational efficiency by causing premature translation termination ^15,16^. The effects of codon usage on elongation kinetics can also feedback to regulate translation initiation process ^7,17,18^.

In addition to directly acting on translation process, accumulating evidence indicates that codon usage influences gene expression by determining mRNA levels genome-wide ^5,6,19–21^.

Codon usage has been shown to indirectly affect translation efficiency by regulating translation-dependent mRNA decay in different organisms ^3,22–26^. In yeast, for example, codon usage influences the recruitment of the CCR4-NOT complex to the ribosome via an interaction between the NOT5 subunit and the ribosomal E-site when translation elongation encounters nonoptimal codons, which can result in a ribosome with an empty A site ^27–29^. The CCR4-NOT complex is the major deadenylating enzyme in eukaryotes and plays a key role in translation-dependent mRNA decay ^30,31^. Thus, the elongation pausing triggered by non-optimal codon promotes mRNA deadenylation and decay. Furthermore, the NOT complex subunits influence elongation kinetics and regulate mRNA solubility in a codon optimality-dependent manner ^31–33^. It is important to note that in addition to its roles in mRNA metabolism, the CCR4-NOT complex is also an important regulator of chromatin structure, transcription, nuclear RNA quality control and mRNA export in the nucleus ^30^, but its nuclear role in the codon usage effect is unknown.

Although effects on translation were thought to be the main mechanism through which codon usage regulates gene expression, recent evidence suggests that codon usage also regulates gene expression by influencing translation-independent processes ^4^. We previously demonstrated that a major effect of codon usage on expression of reporter genes in the filamentous fungus *Neurospora crassa* is through an influence on chromatin structure that impacts transcription efficiency ^5^. Consistent with a translation-independent effect, codon usage or GC content (which correlates with codon usage) within gene coding regions influences mRNA levels without influencing mRNA decay in human cells ^34,35^. Codon usage has also been shown to influence transcription and chromatin structures in *Drosophila* and human cells, suggesting a conserved mechanism in eukaryotes ^21,36^. In addition, non-optimal codons can impact mRNA levels by causing premature transcription termination ^37^. Furthermore, high GC content promotes cytoplasmic mRNA localization, and mRNA splicing enhances nuclear export of AU-rich mRNAs ^38,39^. The nuclear RNA export pathway depends on multiple mRNA export factors such as NXF1, TREX and RBM33 that differentially recognize and facilitate export of either GC-rich or AU-rich transcripts ^19,40–42^. However, it is unclear whether the preferential nuclear export of GC-rich mRNAs is mainly due to the preference at the nuclear export step or at an upstream process. Together, these results suggest that gene codon usage biases are due to selection by both translation-dependent and translation-independent processes.

The relative contributions of translation-independent nuclear effects and translation-dependent effects of codon usage on gene expression in mammalian cells remain unknown. Additionally, the mechanism responsible for the preferential export of GC-rich transcripts, which are enriched with optimal codons, is unclear. In this study, we showed that codon optimality correlates genome-wide with nuclear mRNA and transcription levels in human cells, indicating a broader role for codon usage in regulating nuclear mRNA metabolism. To identify mechanisms involved in codon usage effects in gene expression, we performed a genome-wide CRISPR-Cas9 screen using a dual-codon usage reporter human cell line. This screen identified CNOT4, a RING E3 ligase providing the ubiquitination activity of the CCR4-NOT complex, and many components of the nuclear RNA quality control pathway as factors that mediate the effects of codon usage on gene expression in human cells. Our findings reveal that different nuclear, translation-independent effects of codon usage/nucleotide compositions have important impacts on gene expression levels in human cells by influencing transcription, the nuclear quality control pathway and mRNA localization.

## Results

### Codon usage and nuclear mRNA and transcription levels are correlated genome-wide in human cells

Our previous results on the nuclear role of codon usage in fungi and *Drosophila* cells prompted use to examine the nuclear effect of codon usage genome-wide in human cells. We previously demonstrated in *Drosophila* cells that the genome-wide effect of codon usage on mRNA levels can be masked by tissue-specifically expressed genes and the genome-wide correlations between gene codon usage and mRNA levels are much stronger for constitutively expressed genes than for tissue-specifically expressed genes ^21^. Thus, we compared genome-wide Pearson correlation coefficients between mRNA levels and gene codon usage as measured by codon adaptation indices (CAI) using available total and nuclear RNA-seq data for human cells ^40,43^ for all genes and for constitutively expressed genes. As expected, there was a weak positive correlation between codon usage and total RNA or nuclear RNA levels genome-wide when all genes were included in the analyses (Figure 1a). However, for the constitutively expressed genes (Const1) (i.e., which are up- or downregulated by more than 2-fold in less than five tissues among the 53 human tissues examined) ^21^, the positive correlations became much stronger (Figure 1a). Importantly, the positive correlations were similar for both total and nuclear RNA-seq data. Furthermore, very similar correlation results were seen in two independent global nuclear run-on (GRO-seq) datasets (^44^ and (GSE70449)), which reflect genome-wide transcriptional levels (Figure 1a). Because the nuclear effects of codon usage should be independent of translation, we also performed the analyses of these data using gene GC contents (which reflect nucleotide composition) instead of CAI and found similar positive correlations (Figure 1b). Although CAI and GC content can reflect different aspects of nucleotide composition, CAI values positively correlated with GC contents in the human genome.

**Figure 1.**
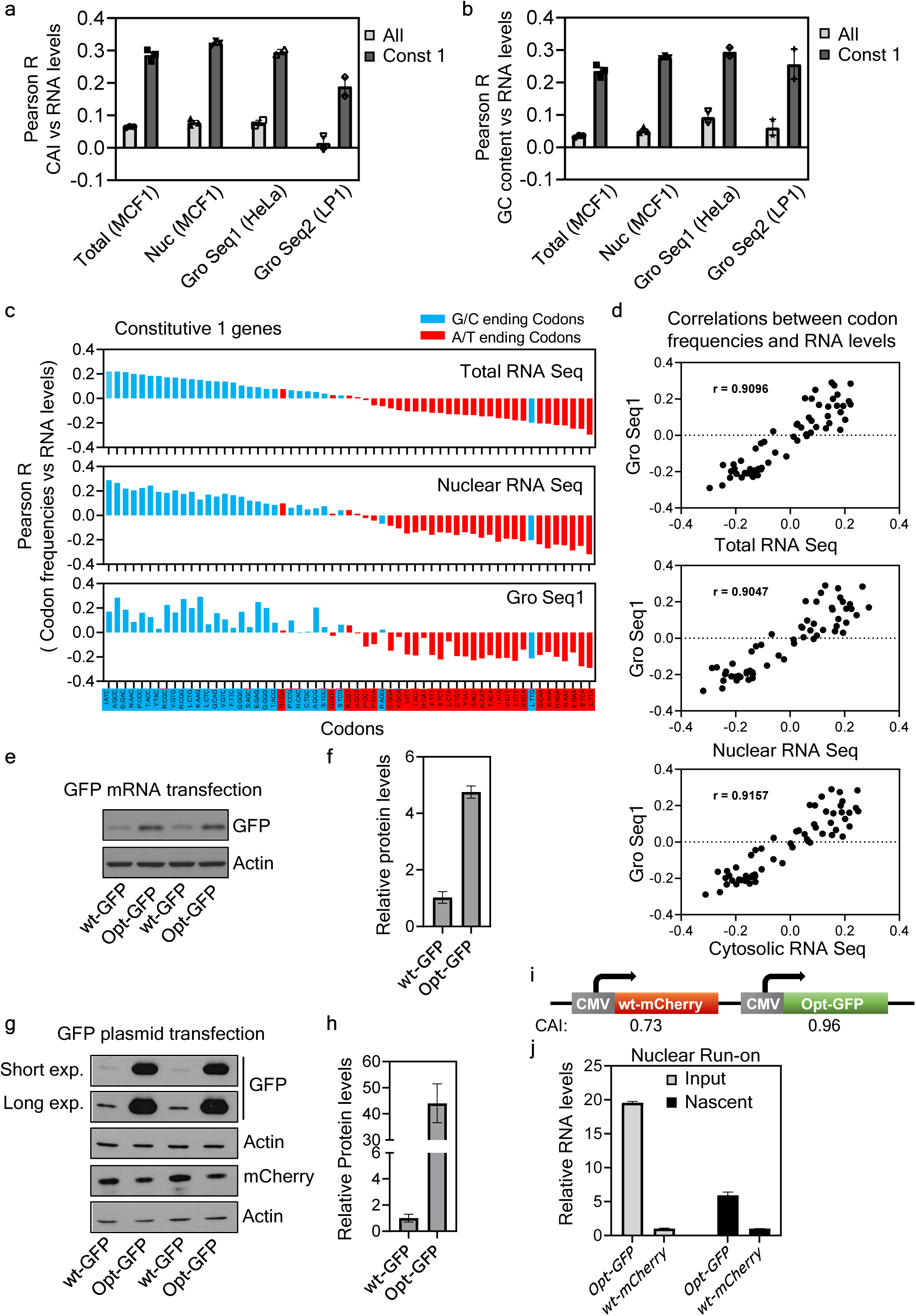
Genome-wide nuclear impact of codon usage on gene expression. (a-b) Plots showing Pearson correlation coefficients between CAI (a) or GC content (b) and abundance of RNA in total, nuclear and nascent RNA samples from human cells as indicated, calculated for either all genes or constitutive1 genes. (c) Pearson correlation coefficients between occurrence of codon frequencies and RNA abundance for 59 synonymous codons in the total, and nuclear RNA samples from MCF1 cells and nascent RNA samples from HeLa cells (Gro Seq) calculated for constitutive1 genes are shown here. The correlation coefficients estimate the effect of codon optimality of each codon on transcript abundance in RNA samples. G/C ending, and A/T ending codons are indicated by blue, and red bars, respectively. (d) Pearson correlation coefficients between occurrence of codon frequencies and RNA abundance for 59 synonymous codons in nascent RNA samples from HeLa cells (Gro Seq) *versus* those in the total, nuclear and cytosolic RNA from MCF1 cells is plotted for constitutive1 genes. Correlation coefficients (r) between different comparisons are indicated. (e) Western blot result showing wild-type (wt) and codon optimized (opt) GFP protein levels in HEK293T cells transfected with indicated *GFP* mRNA. (f) Densitometric analyses of Western blot results of wt and opt-GFP protein levels as in (e) normalized with respective mRNA levels from three independent experiments. (g) Western blot result showing wild-type (wt) and codon optimized (opt) GFP protein levels in HEK293T cells co-transfected with indicated GFP plasmid constructs and opt-mCherry plasmid construct for 72 hrs. mCherry level was used to normalize for transfection efficiency. (h) Densitometric analyses of Western blot results of wt and opt-GFP protein levels as in (g) normalized with mCherry protein levels from three independent experiments. (i) Diagram showing the design of the CMV promoter driven wt-mCherry/opt-GFP dual codon usage reporter construct. CAI of both reporters are indicated. (j) Plot showing relative RNA levels of *opt-GFP* and *wt-mCherry* reporters in nascent and input RNA, measured by RT-qPCR using dual reporter stable cell line. Nascent RNA levels were determined by nuclear run-on assays.

To determine the impact of codon optimality of synonymous codons on mRNA levels, we determined the Pearson correlation coefficients between individual codon frequency (except for stop, methionine, and tryptophan codons) in constitutive genes and their RNA levels using the total mRNA-seq, nuclear RNA-seq, and GRO-seq data. Individual codons were assigned in to G/C ending or A/T ending codon group because human codon usage is biased for C/G ending codons. In all three sets of analyses, the codons that are positively correlated with total, nuclear and Gro-seq RNA levels are almost all G/C ending codons and those with negative correlations are almost all A/T ending codons (Figure 1c). In addition, the codons were also assigned into optimal, intermediate, and non-optimal groups based on a previous study on codon optimality based on RNA stability in human cells ^23^. As shown in Figure S1a, the codons with positive correlations with RNA levels are mostly optimal codons and the codons with negative correlations are mostly non-optimal codons. Furthermore, the correlations of individual codon with RNA levels between the Gro-seq data and those of total RNA, nuclear and cytoplasmic RNA data are highly correlative (Pearson r values > 0.9) (Figure 1d). Together, these results suggest that codon optimality has a genome-wide nuclear effect on gene expression.

To determine if nuclear effect plays a substantial role in determining codon-usage dependent gene expression in human cells, we compared the effect of codon usage on protein expression by transfecting the wild-type and codon-optimized *GFP* mRNAs directly into HEK293T cells (involving mainly translation/cytoplasmic effects) with that by transfecting DNA constructs expressing the same *GFP* genes (involving both nuclear and cytoplasmic effects). As shown in Figure 1e-f, the codon optimization resulted in ∼4-5x increase of GFP protein level (normalized by mRNA levels) by mRNA transfection. In contrast, the codon optimization resulted in more than 40x increase of GFP protein level (normalized with mCherry protein levels from co-transfected *mCherry* DNA construct) by DNA transfection (Figure 1g-h). These results are consistent with our previous results comparing the codon usage effects of the human *Kras* gene (mRNA vs DNA plasmid) ^36^. Together, these results suggest that the nuclear effects of codon usage play a major role in mediating codon-usage dependent gene expression in human cells.

To confirm the transcriptional impact of codon usage, we created a HEK293 cell line stably transfected with a previously created dual-reporter construct that expresses the codon-optimized GFP reporter gene (*opt-GFP*) and the wild-type *mCherry* gene (*wt-mCherry*), which is enriched with rare codons, independently controlled by the same cytomegalovirus (*CMV*) promoter (Figure 1i) ^45^. We then used the dual reporter cell line and nuclear run-on assay to compare the transcription of the two codon usage reporter genes. As shown in Figure 1j, we found that the *opt-GFP* transcription was ∼ 5x as that of the *wt-mCherry*. Thus, codon usage indeed have an important impact on transcription. However, in the input nuclear RNA, we found that the *opt-GFP* mRNA is close to 20x of that of the *wt-mCherry* (Figure 1j). Because the nuclear RNA level is affected by different nuclear effects (transcription, nuclear decay, and export), this result suggests that transcriptional regulation is not the only nuclear process determining the effect of codon usage on mRNA levels. This conclusion is consistent with previous results on nascent RNA production in fungi, *Drosophila*, and human cells when other reporter genes were analyzed ^5,21,35,36,38^. Together, these results suggest that the nuclear, translation-independent effects have an important role in determining how codon usage influences gene expression in eukaryotic organisms.

Because gene GC contents correlate with codon usage bias, the observed codon usage-dependent nuclear nascent RNA effect could be due to mRNA GC content effect. To examine this, we re-analyzed the data in Figure 1c based on the GC contents of individual codons (Figure S1b). Although the 100% GC and 100% AT codons all have positive or negative correlations with nascent RNA levels, respectively, the codons with positive correlations in the 66.7% and 33.3% GC codon groups are almost all optimal codons. These results suggest that the observed codon usage effects are not solely determined by GC contents. Thus, the observed codon usage biases are the selection results by both nucleotide compositions for nuclear effects and optimal translation in cytoplasm.

### A genome-wide CRISPR-Cas9 screen identified many nuclear factors involved in codon usage effect on gene expression

To identify factors involved in regulating the codon usage effect on human gene expression, we created a HEK293 cell line stably transfected with a dual-reporter construct that expresses the codon-optimized mCherry reporter gene (*opt-mCherry*) and the wild-type GFP gene (*wtGFP*), which is enriched with rare codons, independently controlled by *CMV* promoter (Figure 2a) ^45^. As expected, the cell line exhibited much higher mCherry fluorescence levels compared to the levels of GFP fluorescence, indicating a strong codon usage effect on gene expression. We then carried out an unbiased genome-wide CRISPR-Cas9 screen in the opt- mCherry and wtGFP dual reporter cell line by sorting cell population with highest GFP/mCherry fluorescence ratio (0.5% of all cells) and sequencing for enriched single-guide RNAs (sgRNAs) (Figure 2b). This screen allowed the identification of potential factors, when depleted in cells, result in reduced codon usage effect on reporter expression.

**Figure 2.**
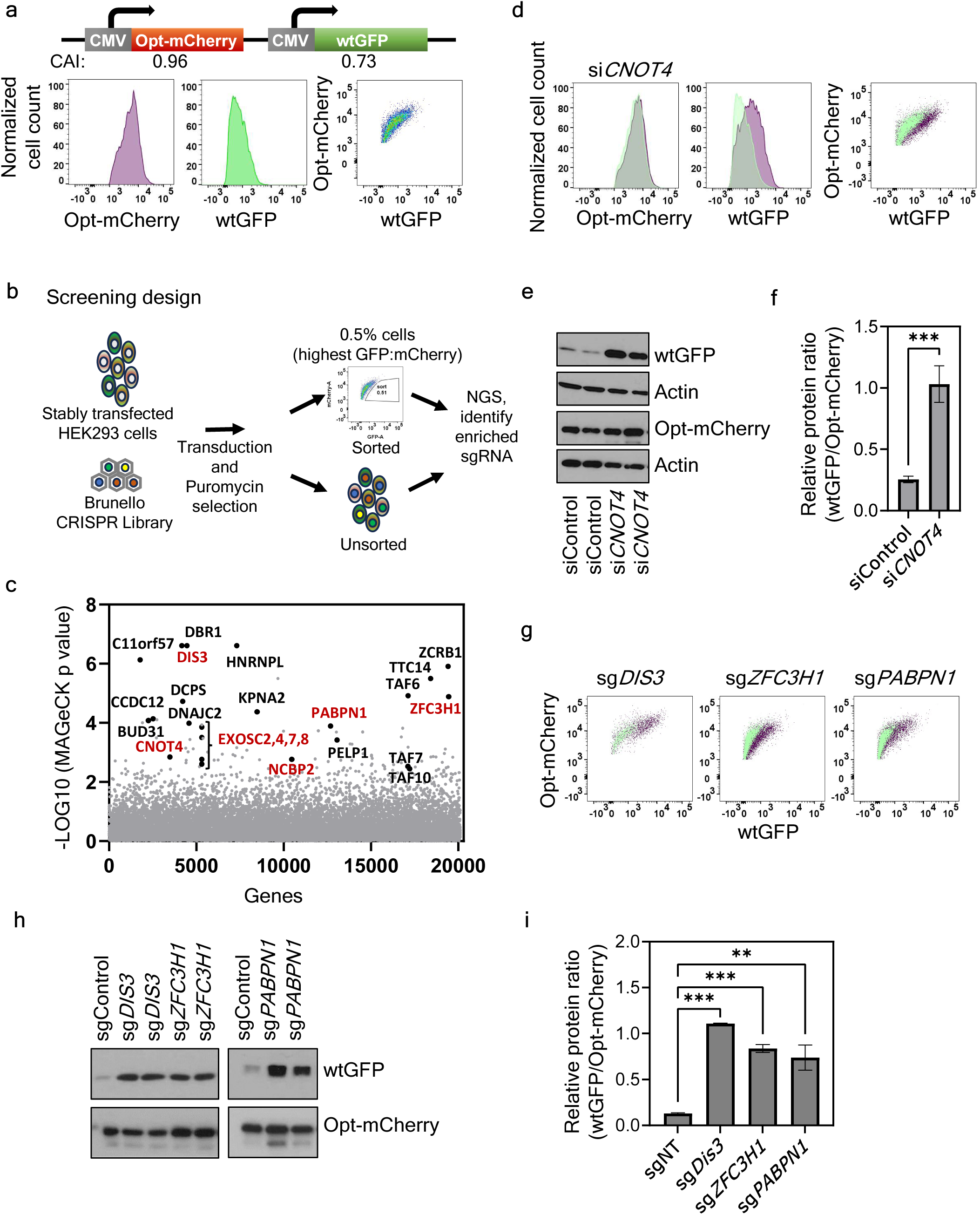
A genome-wide CRISPR–Cas9 screen identified many nuclear factors regulating codon usage effect on gene expression. (a) Diagram showing the design of the CMV promoter driven wtGFP/opt-mCherry dual codon usage reporter construct. CAI of both reporters are indicated. (b) Design of CRISPR–Cas9 screening is depicted here. CRISPR-Cas9 screen was performed using the wtGFP/opt-mCherry dual reporter cell line, cell population with highest GFP/mCherry fluorescence ratio (0.5% of all cells) was sorted by FACS and was used for sequencing to identify enriched sgRNAs. (c) MAGeCK analysis of screening results identified many nuclear factors potentially involved in regulation of codon-usage mediated gene expression (highlighted in black). Genes related to nuclear RNA exosome and CNOT4 are highlighted in red. (d) Validation of screening results by the fluorescence measurements of GFP and mCherry reporters in the wtGFP/opt-mCherry stable cells treated with control siRNA (green) or si*CNOT4* (purple) for 72 h, as assessed by FACS. (e) Western blot results showing wtGFP and opt- mCherry protein levels in the wtGFP/opt-mCherry stable cells treated with control siRNA or si*CNOT4* for 72 h. Actin protein levels are used as loading controls. (f) Densitometric analyses of Western blot results of wtGFP and opt-mCherry protein levels as in (e) from three independent experiments. (g) FACS analysis of wtGFP/opt-mCherry stable cells treated with indicated lentiviral sgRNA and selected with puromycin for 9 days. Non targeting control sgRNA treated cells are shown in green while knockout cells are shown in purple in dot plots. (h) Western blot result showing wtGFP and opt-mCherry protein levels in the wtGFP/opt-mCherry stable cells following sgRNA treatment as indicated. (i) Densitometric analyses of Western blot results of wtGFP and opt-mCherry protein levels as in (h) from three independent experiments. Data in (f) and (i) are presented as mean ±SEM, one-way ANOVA with Dunnett’s multiple comparisons test, **P < 0.01; ***P < 0.001.

The top hits of the screen include CNOT4, a subunit of the CCR4-NOT complex, in addition to many factors localized to the nucleus (Figure 2c). These factors include the multiple RNA exosome complex components EXOSC2, 4, 7 and 8; the catalytic subunit of nuclear RNA exosome DIS3; components of the nuclear PAXT complex, which binds the poly-A tails of nascent mRNAs and target them to the nuclear exosome for decay ^46^, including the zinc-finger protein ZFC3H1, the nuclear poly-A binding protein PABPN1, and the nuclear CAP binding protein NCBP2 (also known as CBP20); transcription factors TAF6, 7, 10; and other proteins known to be enriched in the nucleus. Consistent with a previous study on the role of codon usage-dependent mRNA splicing and export ^38^, several nuclear factors involved in splicing (DBR1, ZCRB1, and HNRNPL) were also identified as top hits in the screen.

Because the CCR4-NOT complex was previously shown to regulate translation-dependent non-optimal codons-mediated mRNA decay and mRNA solubility in yeast ^27,31,33^, the identification of CNOT4 suggest that it also plays a role in regulating the codon usage-dependent effects on gene expression in human cells. When the *wtGFP/opt-mCherry* dual reporter cells were depleted of CNOT4 using a small interfering RNA (siRNA) targeting *CNOT4*, there was a significant increase in wtGFP fluorescence levels but not opt-mCherry (Figure 2d). In addition, there was a specific increase of wtGFP protein level (Figure 2e-f). To confirm the codon usage-dependent effect of CNOT4 depletion, we also examined the *opt-GFP/wt-mCherry* dual-reporter cells to determine if the codon usage effect observed is reporter-specific. As shown in Figure S2a-c), there was a specific increase of wt-mCherry protein levels compared to those of opt-GFP after CNOT4 depletion, indicating that the effects observed are codon usage-specific and are independent of reporter genes used.

The identification of DIS3 and components of the PAXT complex suggested that the nuclear exosome and the nuclear RNA quality control are involved in mediating the codon usage effect. PAXT complex directs polyadenylated RNA substrates to the nuclear exosome core for decay ^46^. To confirm their role, we separately depleted DIS3, ZFC3H1, and PABPN1 in the *wtGFP/opt-mCherry* dual reporter cells by using targeted sgRNAs. As shown in Figure 2g and 2h-i, depletion of each of these proteins led to a significant and specific increase of wtGFP fluorescence and GFP protein levels without significant effects on opt-mCherry fluorescence or opt-mCherry protein levels. These results confirmed the involvement of these factors in mediating codon usage effect on gene expression.

### CNOT4 impacts codon usage-dependent gene expression largely due to its nuclear effect on mRNA levels

To determine whether the codon usage effect of CNOT4 is mainly mediated by the translation-dependent role of the CCR4-NOT complex, we transfected 5’ capped and 3’ polyadenylated mCherry mRNA that was either codon-optimized or not into HEK293 cells. The transfection of the mRNA instead of plasmid DNA eliminated potential transcription-dependent nuclear effects that can influence cytoplasmic mRNA levels. Surprisingly, depletion of CNOT4 did not significantly impact the codon usage effect on mCherry protein levels (normalized with transfected mRNA levels) (Figures 3a-b), suggesting that the translation-dependent effect on translation efficiency or mRNA decay is not the main mechanism that determines the codon usage effect of CNOT4 for the reporter transgenes.

**Figure 3.**
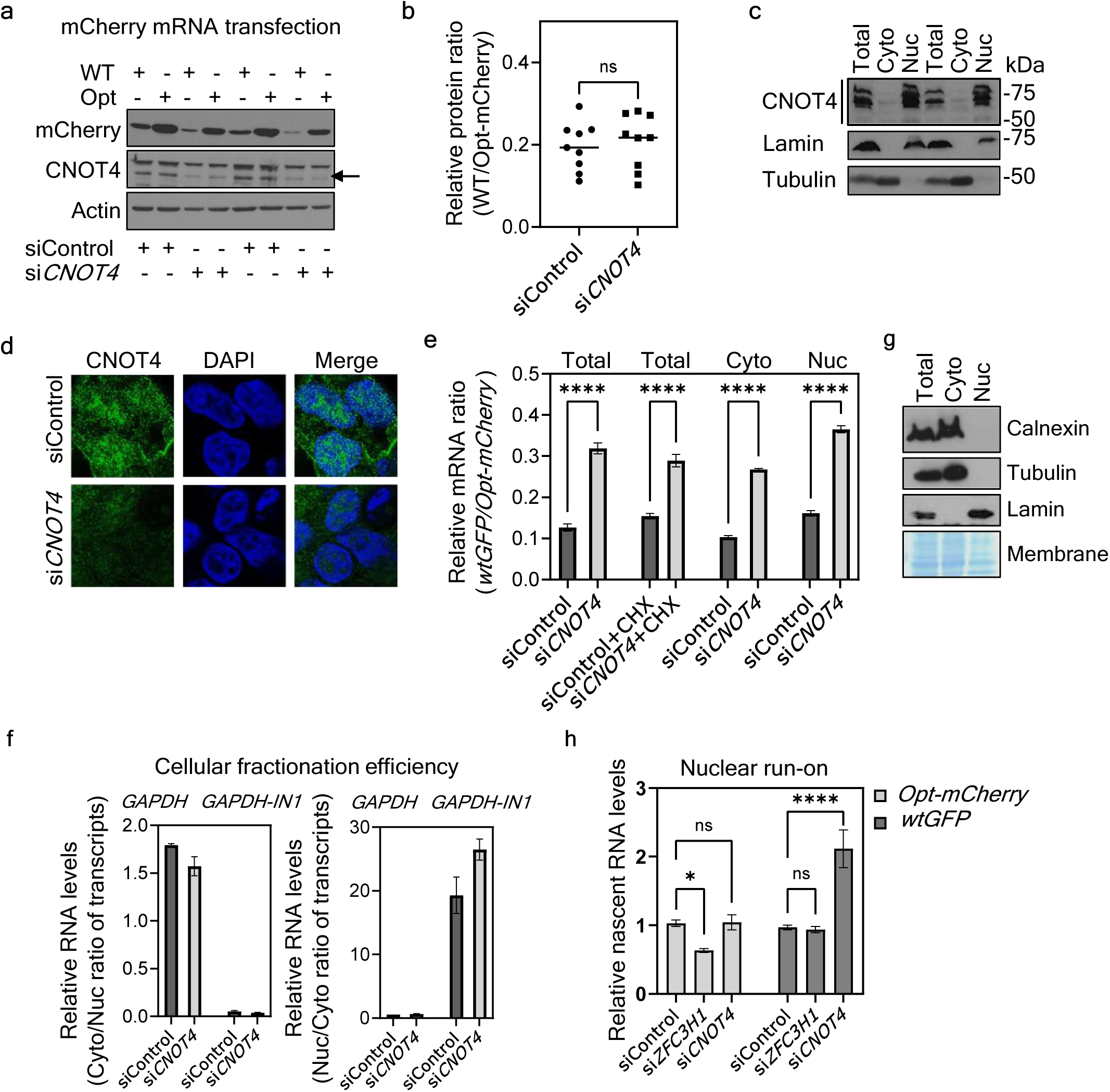
CNOT4 impacts codon usage effects on reporter gene expression in human cells largely by its nuclear effect on mRNA levels (a) Western blot result showing wild-type (WT) and optimal (Opt) codons enriched mCherry protein levels in HEK293 cells first treated with siControl or si*CNOT4* for 72 h and then transfected with indicated *mCherry* mRNA for 8 h. Western blot results confirmed depletion of CNOT4 protein after siRNA mediated knockdown (middle panel). Actin protein levels are used as loading controls. (b) Densitometric analyses of Western blot results of WT and Opt-mCherry protein levels as in (a) normalized with mRNA levels from three independent experiments. ns - non significant. (c) Western blot results showing indicated protein levels in total, cytosolic and nuclear protein fractions of HEK293 cells. (d) Representative immunofluorescence imaging results in HEK293 cells treated with siControl or si*CNOT4* for 72 h using a rabbit polyclonal anti-CNOT4 antibody. (e) Quantification of the ratio of mRNA levels of *GFP* and *mCherry* reporters in total, cytosolic and nuclear RNA fractions isolated from the wtGFP/opt-mCherry stable cells treated with indicated siRNA for 72 h and with cycloheximide (CHX) as indicated for 24 h as measured by RT-qPCR. Data are presented as mean ±SEM, n=3, one-way ANOVA with Sidak’s multiple comparisons test, ****P < 0.0001. (f) Quantification of the ratio of cytoplasmic to nuclear levels (left panel) or nuclear to cytoplasmic levels (right panel) of *GAPDH* mRNA and *GAPDH-IN1* (intron1 containing) pre- mRNA in RNA samples from Figure 1e as measured by RT-qPCR to determine cellular fractionation efficiency. (g) Western blot results showing indicated protein levels in total, cytosolic and nuclear protein fractions of HEK293 cells. Coomassie blue stained membrane is shown below to estimate protein loading. (h) Nuclear run-on assay results showing relative nascent *opt-mCherry* and *wtGFP* RNA levels (normalized with *18S* RNA levels) as measured by RT-qPCR using the wtGFP/opt-mCherry stable cells. The cells were subjected to indicated siRNA mediated knockdown for 72 h. Data are presented as mean ±SEM, n=3, unpaired two- tailed Mann-Whitney test, ns - non significant; *P < 0.05; ****P < 0.0001.

The NOT proteins were originally identified as global transcriptional regulators and are known to participate in various steps of transcriptional process in yeast ^30,31,47^. To evaluate whether CNOT4 has a function in the nucleus of human cells, we examined its cellular localization in HEK293 cells. Western blot analysis showed that various CNOT4 isoforms were highly enriched in the nuclear fraction and their cytoplasmic levels were low (Figure 3c), suggesting that the CNOT4 mainly acts in the nucleus. Immunofluorescence assay also confirmed the nuclear enrichment of CNOT4 in HEK293 cells although some signals were also seen in the cytoplasm (Figure 3d). The immunofluorescence signals were reduced substantially in the cells depleted of CNOT4, confirming the specificity of the fluorescence signal for CNOT4. Consistent with these results, it was previously shown that CNOT1 and CNOT3, two other subunits of the CCR4-NOT complex, were also present in the nuclei of human and mouse cells48,49.

The nuclear enrichment of CNOT4 prompted us to determine whether the codon usage effect we observed is largely caused by the nuclear effect of CNOT4. We compared the ratios of *wtGFP* mRNA to *opt-mCherry* mRNA in the total, cytoplasmic, and nuclear RNA preparations of the dual reporter cells. The *opt-mCherry* mRNA was detected at significantly higher levels than the *wtGFP* mRNA in total RNA samples of the control cells (Figure 3e), indicating that the codon usage effect is largely due to changes in mRNA levels. *CNOT4* depletion not only resulted in a marked increase in the *wtGFP*/*opt-mCherry* mRNA ratios in the total and cytoplasmic RNA samples but also in nuclear RNA (Figure 3e). Importantly, a marked increase in the *wtGFP*/*opt- mCherry* mRNA ratios was observed in the total RNA in the *CNOT4* depleted cells compared to control cells even in the presence of a translation inhibitor, cycloheximide, suggested that this effect of CNOT4 on the reporter mRNA levels is largely translation independent (Figure 3e).

Relative *GAPDH* mRNA and nascent intron-containing *GAPDH* pre-mRNA levels in the cytosolic and nuclear RNA fractions demonstrated the purity of our cellular fractionation preparations (Figure 3f). To exclude the possibility of contamination of mRNA attached to ER membrane in nuclear fractions, we examined the levels of Calnexin, an ER marker, in total, cytoplasmic and nuclear fractions. As seen in Figure 3g, Calnexin was present in total and cytoplasmic fractions but not in the nuclear fraction. In addition, tubulin was found in the total and cytoplasmic fractions but not in nuclear fraction. On the other hand, the nuclear marker Lamin was absent from the cytoplasmic fraction. These results further confirmed the purity of our cytoplasmic and nuclear fractionations. These results suggest that CNOT4 regulates codon usage effect on gene expression by preferentially suppressing the nuclear levels of mRNAs enriched in non-optimal codons.

We then examined if CNOT4 affects the codon-usage dependent transcription of reporter genes by nuclear run-on assays. As shown in Figure 3h, CNOT4 silencing indeed resulted in a significant increase of *wtGFP* transcription while no effect was seen at the transcription levels of *opt-mCherry* as compared to control cells, suggesting a role of CNOT4 in mediating codon usage-dependent transcriptional regulation. This result is consistent with the known role of CCR4-NOT complex as a transcriptional regulator ^30^. On the other hand, the depletion of the PAXT component ZFC3H1 did not affect the transcription of *wtGFP* as compared to control cells (Figure 3h). Thus, CNOT4 and PAXT complex have different roles in the nuclear effects of codon usage.

### Depletion of CNOT4 impairs genome-wide codon usage effects

We next examined the global effect of CNOT4 depletion on RNA levels genome-wide by sequencing RNA from total and nuclear fractions of HEK293 cells treated with an siRNA specific for *CNOT4*. Analyses of the total RNA-seq data revealed that CNOT4 depletion resulted in a significant reduction of correlation between CAI or GC content and mRNA levels (Figure 4a and S3). In addition, the fold changes of mRNA levels in CNOT4 depleted cells as compared to control treated cells showed clear negative correlations for both CAI and GC contents (Figure 4b-c). Upregulated mRNAs are preferentially genes with non-optimal codon usage profiles, which is consistent with its role in transcription by suppressing genes enriched with non-optimal codons. Furthermore, depletion of CNOT4 resulted in downregulation of nuclear mRNAs enriched for genes involved in chromatin regulation (Figure 4d), which is consistent with the known role of the CCR4-NOT complex in transcriptional regulation. This conclusion is also consistent with our previous studies that demonstrated the roles of chromatin regulators in codon usage-dependent transcriptional effects in other organisms ^5,36,50^.

**Figure 4.**
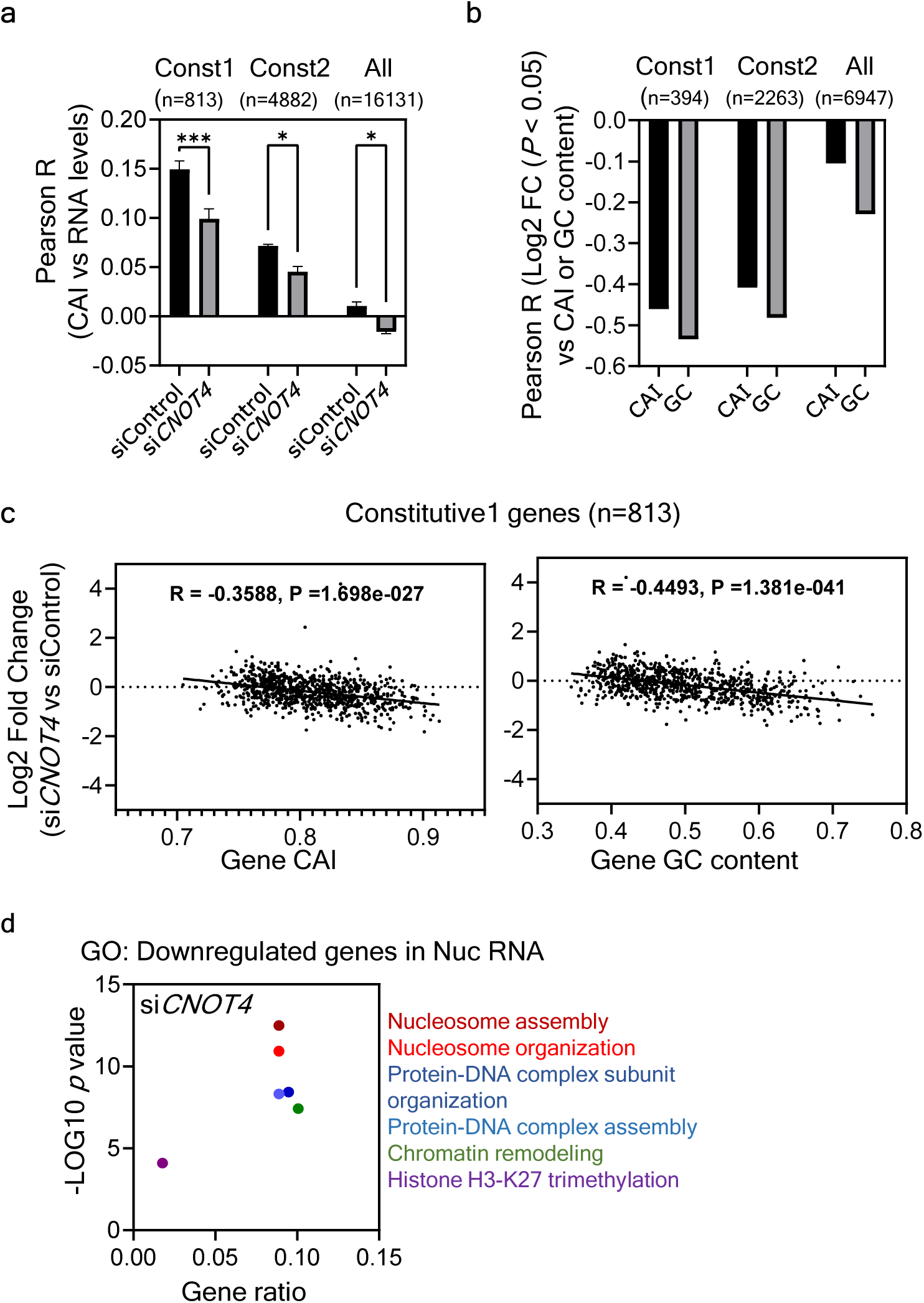
Genome-wide effects of CNOT4 depletion on codon usage-mediated mRNA levels. (a) Plots showing Pearson correlation coefficients between CAI (codon-usage mediated effects) and abundance of RNA in total RNA samples from HEK293 cells treated with either siControl or si*CNOT4* as indicated for 72 h followed by RNA sequencing, calculated for all genes or constitutive genes. Data is presented as mean ±SEM, n=2, one-way ANOVA with Sidak’s multiple comparisons test, *P < 0.05; ***P < 0.001. (b) Plot showing Pearson correlation coefficients between Log2 fold change in total RNA levels in CNOT4 depleted cells as compared to control treated cells and their CAI or GC content. (c) Scatter plots showing Pearson correlation coefficients between Log2 fold change in RNA levels (in CNOT4 depleted cells as compared to control treated cells) vs their CAI (top) or GC content (bottom) of constitutive1 genes. Spearman’s R and *P* values are indicated. Dots, genes. (d) Gene ontology (GO) enrichment analysis of the downregulated genes in nuclear RNA of CNOT4 depleted cells compared to control cells.

### The nuclear, but not cytosolic, RNA exosome and the nuclear PAXT complex mediate the codon usage effect on gene expression

The multi-subunit RNA exosome is located in both nucleus and cytosol of eukaryotic cells ^51^ (Figure 5a). In the nucleus, its primary function is in RNA quality control as the RNA exosome processes and degrades a variety of aberrant noncoding and pre-mRNA transcripts. In the cytoplasm, the complex is involved in mRNA turnover and quality control. The CCR4-NOT complex is known to mediate mRNA decay in a translation- and codon usage-dependent manner by monitoring the speed of translation elongation and causing mRNA deadenylation and subsequent decay ^22,23,27,29^. DIS3L is the catalytic subunit of the cytosolic exosome ^52^. Its co- factor is the superkiller (Ski) complex, which consists of SKIV2L, the tetratricopeptide repeat-containing protein TTC37 (also known as Ski3), and two copies of the WD40-repeat protein Ski8 ^53^. In the cytosol, HBS1L (also known as Ski7) connects the Ski complex to the RNA exosome for co-translational mRNA quality control ^54^.

**Figure 5.**
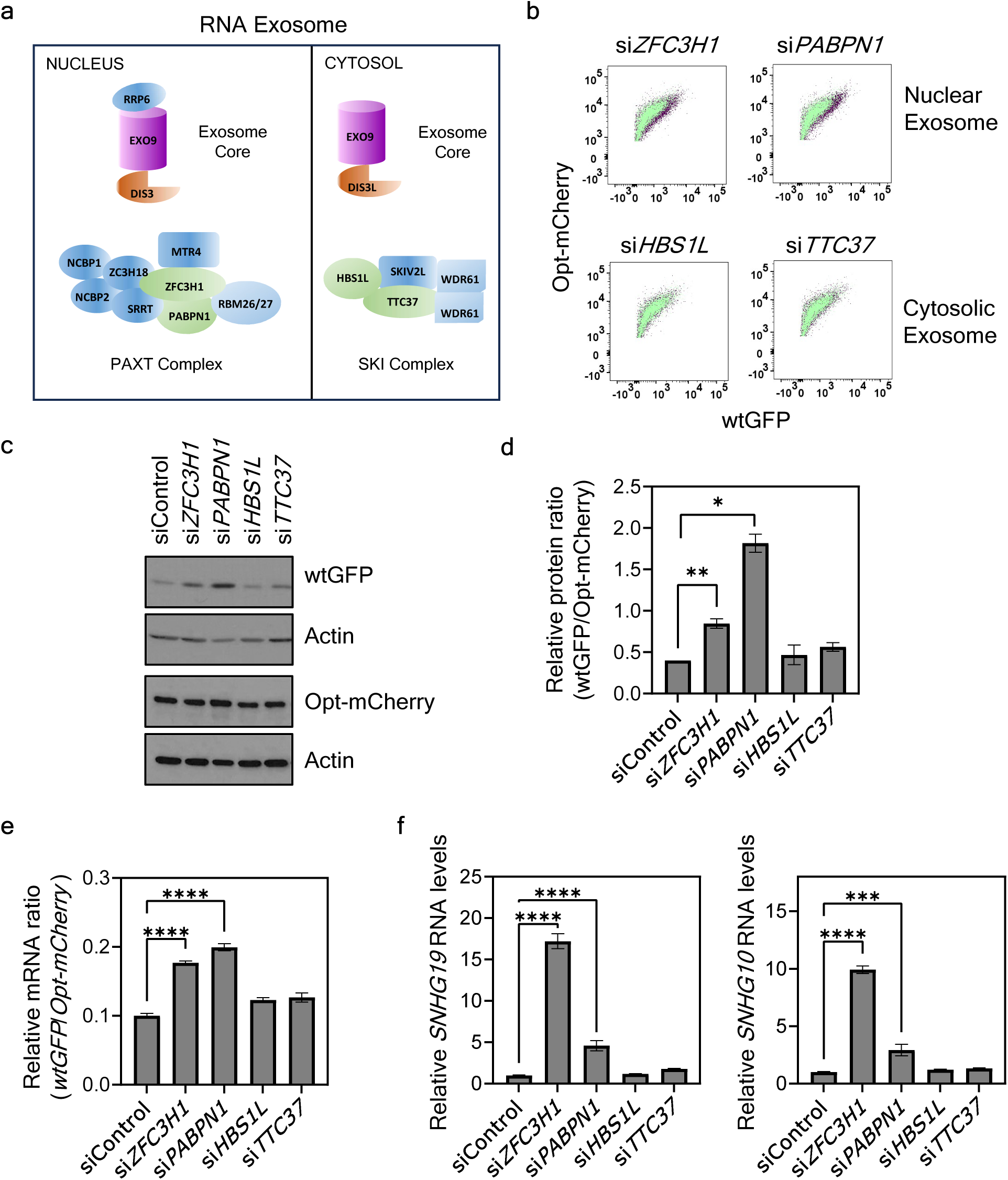
Nuclear but not cytosolic RNA exosome components participate in mediating the codon usage effect on gene expression. (a) A schematic showing components of nuclear and cytosolic RNA exosome and their RNA targeting complexes. (b) FACS analysis of wtGFP/opt- mCherry stable cells following siRNA mediated knockdown of indicated genes. Control siRNA treated cells are shown in green while knockdown cells depleted of different factors are shown in purple in dot plots. (c) Western blot result showing wtGFP and opt-mCherry protein levels in wtGFP/opt-mCherry stable cells following siRNA mediated knockdown of genes as indicated. Actin protein levels are used as loading controls. (d) Densitometric analyses of Western blot results of wtGFP and opt-mCherry protein levels as in (c) from three independent experiments. (e) Quantification of the ratio of mRNA levels of *GFP* and *mCherry* in total RNA isolated from wtGFP/opt-mCherry stable cells treated with indicated siRNA for 72 h as measured by RT-qPCR. (f) Quantification of *SNHG19* (left) and *SNHG10* (right) RNA levels in total RNA isolated as in (e). Data in (d-f) are presented as mean ±SEM, n=3, one-way ANOVA with Dunnett’s multiple comparisons test, *P < 0.05; **P < 0.01; ***P < 0.001; ****P < 0.0001.

To identify the exosome and adaptor complex that regulate codon usage-dependent expression of our reporter genes, we used siRNA to silence expression of different components of these complexes. siRNA-mediated depletions of PAXT complex components ZFC3H1 and PABPN1 significantly increased the expression of wtGFP but not opt-mCherry in the wtGFP/opt-mCherry dual reporter cell line, whereas depletion of DIS3L, HBS1L, or TTC37 did not significantly affect the expression of reporter proteins despite of the accumulation of previously identified cytoplasmic exosome substrates ^55^ (Figure 5b-d and S4a). We also investigated the codon usage dependent effect on the expression of our reporter genes at the transcripts level and found a significant increase in *wtGFP*/*opt-mCherry* mRNA ratio in the cells depleted of ZFC3H1 and PABPN1, but not in HBS1L and TTC37 depleted cells (Figure 5e). In addition, we examined at the levels of previously identified PAXT complex substrates, *SNHG19* and *SNHG10* ^46^, and found that depletions of ZFC3H1 and PABPN1 result in an accumulation of *SNHG19* and *SNHG10* RNA, while HBS1L and TTC37 depleted cells did not show any effect on these substrates (Figure 5f), confirming that PAXT and Ski complexes are functionally distinct. Overall, these results suggest that the nuclear RNA exosome and the PAXT complex but not the cytosolic RNA exosome components mediate the observed codon usage effect on reporter gene expression.

Two RNA-binding proteins, RBM26 and RBM27, were recently identified as components of the nuclear PAXT complex; loss of either protein results in an accumulation of some RNA substrates of the PAXT complex ^56^. RBM26 and RBM27 co-depletions resulted in a dramatic accumulation of the PAXT substrate *SNHG19* RNA but did not alter the expression of reporter proteins in a codon usage-dependent fashion (Figure S4b-c), suggesting that the PAXT complex mediates codon usage-dependent nuclear mRNA decay independently of RBM26 and RBM27. Thus, the RBM26 and RBM27-associated PAXT may specifically target aberrant noncoding and pre-mRNA transcripts but not mature mRNAs of non-optimal codon usage.

### PAXT complex affects codon usage-dependent nuclear mRNA decay

ZFC3H1 acts as a nuclear retention factor for polyadenylated RNA substrates by competing with the RNA export factors ^57^. Similarly, PABPN1 prevents nuclear export of unspliced RNA ^58^. We therefore investigated if these PAXT components affect the relative reporter mRNA levels in a codon usage-dependent fashion in the total, cytoplasmic, and nuclear RNA preparations. As shown in Figure 6a, siRNA-mediated silencing of *ZFC3H1* and *PABPN1* resulted in an increase in the ratio of *wtGFP*/*mCherry* mRNA levels in both nuclear and cytoplasmic RNA fractions. The specific presence of Tubulin and Lamin in the cytoplasmic and nuclear preparations respectively, indicated the success of our fractionation procedure (Figure 6b). As expected, we observed an increase in wtGFP amounts in total protein and cytosolic protein fractions in ZFC3H1 and PABPN1 depleted cells as compared to control treatment, whereas the levels of opt-mCherry protein remained unchanged (Figure 6b). The RNA fractionation efficiency was confirmed by the enrichment of *GAPDH* mRNAs in cytosolic fraction and of intron-containing *GAPDH* pre-mRNAs in the nuclear RNA fractions (Figure S5a-b). The efficiency of ZFC3H1 and PABPN1 depletion was confirmed by the accumulation of *SNHG19* RNA (Figure S5c). Thus, loss of PAXT activity results in an increased export of *wtGFP* mRNA to cytoplasm, suggesting that the PAXT acts to prevent the export of mRNAs enriched with rare codons in the nucleus.

**Figure 6.**
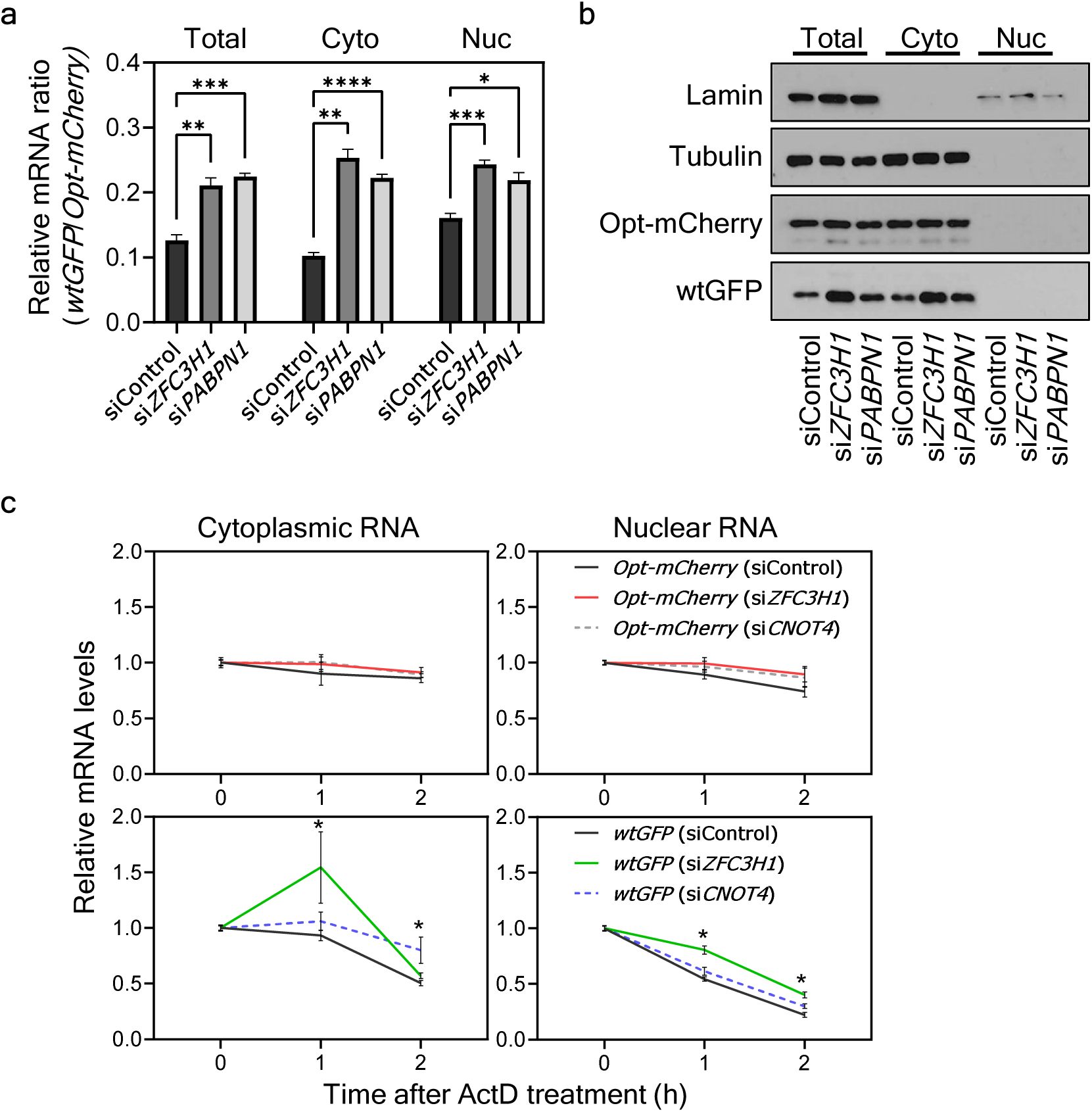
PAXT complex affects codon usage-dependent nuclear mRNA decay (a) Quantification of the ratio of mRNA levels of *GFP* and *mCherry* reporters in total, cytosolic and nuclear RNA fractions isolated from wtGFP/opt-mCherry stable cells treated with indicated siRNAs for 72 h as measured by RT-qPCR. Data are presented as mean ±SEM, n=3, one-way ANOVA with Dunnett’s multiple comparisons test, *P < 0.05; **P < 0.01; ***P < 0.001; ****P < 0.0001. (b) Western blot result showing wtGFP and opt-mCherry protein levels in total, cytosolic and nuclear protein fractions isolated from wtGFP/opt-mCherry stable cells as in (a). Lamin and Tubulin protein levels are used as controls for subcellular fractionation efficiency. (c) Quantification of relative mRNA levels of *GFP* and *mCherry* reporters in cytoplasmic and nuclear RNA fractions isolated from wtGFP/opt-mCherry stable cells subjected to siRNA mediated knockdown for 72 h followed by actinomycin D treatment for indicated time intervals. RNA levels were normalized with *GAPDH* mRNA levels in respective samples, as measured by RT-qPCR. Data are presented as mean ±SEM, n=3, unpaired two-tailed Mann-Whitney test, *P < 0.05.

Although codon optimality is known to affect translation-dependent mRNA decay, whether it affects mRNA decay or clearance in the nucleus is unknown. Thus, we examined the decay rates of *wtGFP* and *opt-mCherry* mRNAs in the nucleus after the addition of transcription inhibitor actinomycin D. In the presence of the transcription inhibitor, the *wtGFP* mRNA was cleared from the nucleus much faster than the *opt-mCherry* mRNA (Figure 6c, comparing top and bottom panels). Because mRNA export is known to be affected by codon usage and GC-rich mRNAs are preferentially exported ^38,40^, this result suggests that the rapid clearance of *wtGFP* mRNA may be due to its rapid decay in the nucleus rather than its preferential export. Although ZFC3H1 depletion did not affect the *opt-mCherry* mRNA clearance rate from the nucleus, it significantly slowed *wtGFP* mRNA clearance (Figure 6c, right panels), suggesting that PAXT complex preferentially mediates the clearance of mRNAs enriched with non-optimal codons.

Thus, in addition to removing aberrant transcripts in the nucleus, the nuclear RNA quality control machinery also preferentially targets mRNAs with non-optimal codon usage for decay through the nuclear exosome. In the cytoplasmic fractions also, depletion of ZFC3H1 resulted in a significant increase in *wtGFP* but not *opt-mCherry* transcripts levels (Figure 6c, left panels), suggesting that *wtGFP* mRNA stabilization in nucleus may be coupled with an increased export to cytoplasm.

We also examined whether CNOT4 is involved in mRNA decay in the human cell nucleus. CNOT4 depletion did not result in a significant change of *wtGFP* transcript decay in the nucleus, while an increase in *wtGFP* transcript levels was observed in cytoplasmic RNA after 2 hrs of actinomycin D treatment (Figure 6c, lower panels). This latter result is consistent with a role for the CCR4-NOT complex in promoting the decay of mRNA enriched with non-optimal codons in cytoplasm ^27,29,31^. Unlike ZFC3H1 depletion, CNOT4 depletion did not increase *SNHG19* RNA accumulation (Figure S5d), indicating that CNOT4 does not act through the PAXT complex. Thus, CNOT4 and PAXT play different roles in the nuclear codon usage effects: CNOT4 mainly suppresses transcription of genes enriched with non-optimal codons while PAXT promotes the nuclear decay of RNA enriched with non-optimal codons. Together, these combined effects result in preferential cytoplasmic accumulation of mRNAs enriched with optimal codons.

### PAXT components affect global codon usage-dependent mRNA cellular localization

To determine the genome-wide effects of PAXT components on gene expression, we depleted ZFC3H1 and PABPN1 from HEK293 cells using siRNA and then sequenced RNAs from total, cytoplasmic, and nuclear fractions. To determine the mRNA cellular distribution, we calculated the ratio between cytoplasmic and nuclear mRNA levels for each gene. For constitutively expressed gene groups (constitutive 1: up- or downregulated by >2-fold in no more than five human tissues; constitutive 2: up- or downregulated by >3-fold in no more than seven tissues) ^21^, there are positive correlations between mRNA location and CAI/GC contents (Figure 7a-b), indicating that optimal codon usage promotes preferential cytoplasmic distribution of mRNAs with optimal codon usage. Depletion of either ZFC3H1 or PABPN1 resulted in a dramatic decrease in the positive correlation between gene CAI/GC content and ratios of cytoplasmic/nuclear RNA levels for constitutively expressed genes (Figure 7a-b), indicating a key role for the PAXT components in the regulation of codon usage-dependent mRNA localization. When the analyses were performed for total RNA levels instead of the cytoplasmic/nuclear RNA ratios, depletion of ZFC3H1 or PABPN1, only resulted in a slight decrease in the Pearson R correlation between CAI and RNA levels of the constitutive genes as compared to control cells (Figure S6a), suggesting that the effect of PAXT complex on codon-usage dependent gene expression are mainly mediated by influencing the mRNA localization in the cells.

**Figure 7.**
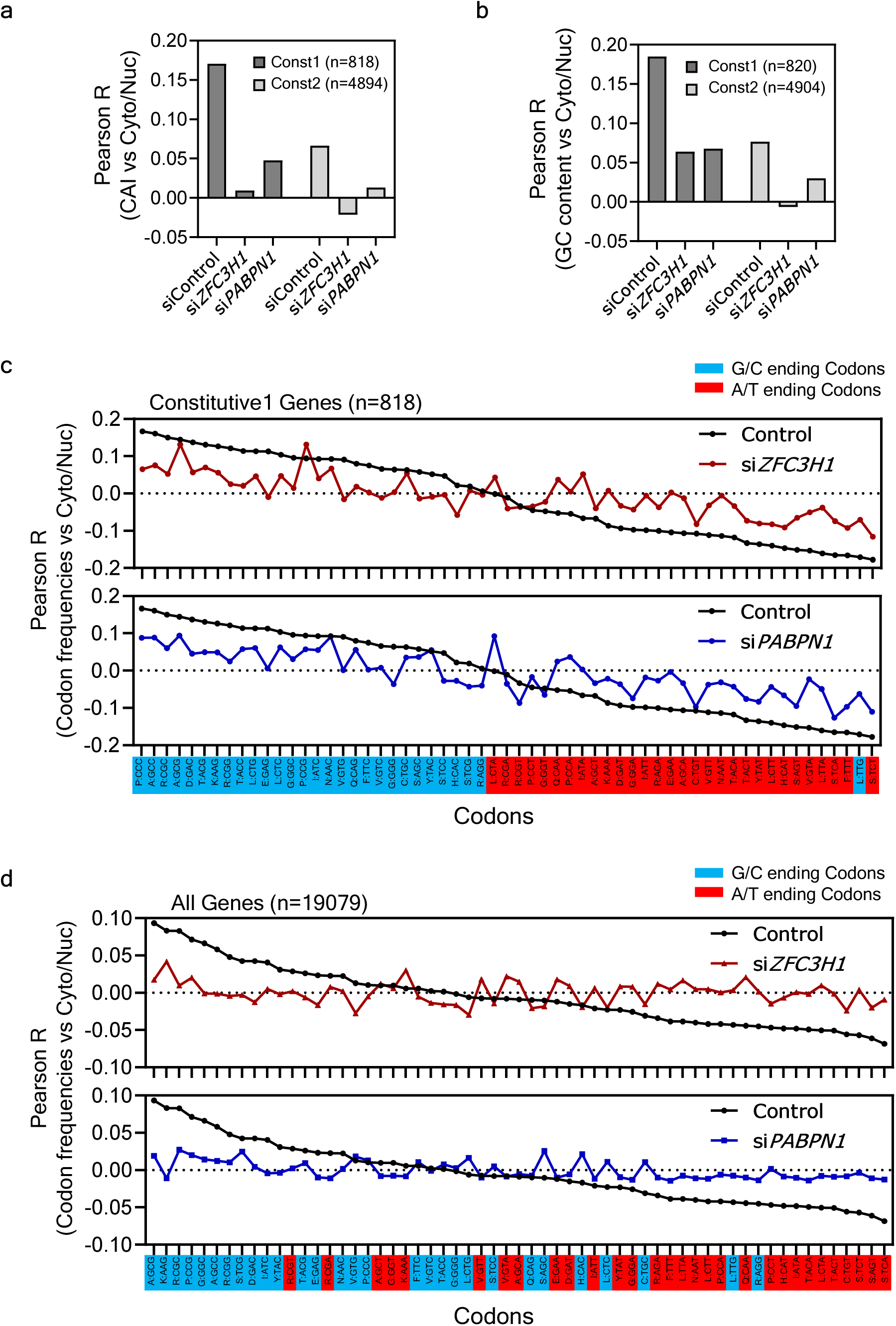
PAXT complex components mediate genome-wide codon usage effects on mRNA localization. (a) Plot showing Pearson correlation coefficients between CAI (to evaluate the codon-usage mediated effects) and cytoplasmic/nuclear ratio of mRNA (a measure of cytoplasmic localization of RNA) in HEK293 cells treated with indicated siRNA for 72 h followed by cellular fractionation and RNA sequencing, calculated for constitutive genes as indicated. (b) Plot showing Pearson correlation coefficients between GC content (a measure of nucleotide composition) and cytoplasmic/nuclear ratio of mRNA in ZFC3H1 and PABPN1 knockdown cells, calculated for constitutive genes as indicated. (c-d) Pearson correlation coefficients between occurrence of codon frequencies and cytoplasmic/nuclear ratio of mRNA for 59 synonymous codons in RNA samples from the control cells and the ZFC3H1 and PABPN1 knockdown cells for constitutive1 (c) or all genes (d) to estimate the effect of optimality of each codon on transcript abundance. G/C ending, and A/T ending codons are indicated by blue, and red bars, respectively. Number of genes (n) used in analyses in different groups are shown in parenthesis.

To determine the effect of individual codons on mRNA localization, Pearson correlations between gene codon frequencies and cytoplasmic/nuclear ratios were determined. When only constitutive genes were used in the analysis, codons that were positively correlated with mRNA cytoplasmic/nuclear ratios are all G/C ending codons whereas codons with negative correlations were almost all A/T ending codons (Figure 7c). A similar observation was also observed when synonymous codons were assigned as optimal codons and non-optimal as previously described for mRNA stability in human cells ^23^ (Figure S6b). Most of the exceptions in the latter analysis are codons that end in G/C nucleotide (and are not the rarest synonymous codons). These results further confirm the genome-wide codon usage effect on mRNA localization. It is important to note that a similar codon optimality effect is also seen when all expressed genes were included in the analysis (Figure 7d and S6c), indicating that the codon usage effect on mRNA localization is global and not limited to constitutively expressed genes.

When either ZFC3H1 or PABPN1 was depleted, the codon usage-dependent correlations for both G/C ending/optimal and A/U ending/non-optimal codons were markedly affected for almost all codons (Figure 7c-d and S6b-c, red or blue data points), indicating global impairment of the codon-usage-dependent mRNA localization in the absence of these PAXT components. It should be noted that the analyses using constitutive genes (Figure 7c and S6b) and all genes (Figure 7d and S6c) resulted in similar observations, indicating that the effect of PAXT on codon usage-dependent mRNA localization is broad and not specific for constitutive genes.

## Discussion

Codon usage was previously thought to mediate its effect on gene expression mainly through translation-dependent processes such as translation elongation, initiation, premature termination, and translation elongation-dependent mRNA decay ^4,29,31^. In this study, we showed that translation-independent nuclear effects of codon usage also play a major role in determining gene expression levels by affecting different nuclear gene regulatory processes, including transcription and nuclear RNA quality control. Our results suggest that the evolution of codon usage biases is due to selection on both translation-related processes in the cytoplasm and nuclear translation-independent processes. For the latter, the open-reading frame sequences are not read as codons but nucleotide compositions that resemble codon usage biases. The nucleotide compositions are recognized by the nuclear machineries in forms of DNA/RNA elements to activate or suppressing transcription or nascent RNA stability/export.

By examining the relationship between codon usage and nuclear RNA level/transcription level, we showed that codon optimality correlates genome-wide with nuclear mRNA and transcription levels in human cells, suggesting a broader role for codon usage/nucleotide composition in regulating nuclear mRNA levels. Individual genome-wide codon optimality for total mRNAs and nuclear nascent mRNAs are highly similar to each other, indicating that selection on transcription also results in codon usage/GC compositions similar to those selected by translation-dependent processes.

To identify factors involved in regulating codon usage effects on gene expression, we performed a non-biased genome-wide CRISPR-Cas9 screen in a dual codon usage reporter cell line. The identified factors include the CCR4-NOT subunit CNOT4 and many factors involved in nuclear RNA quality control pathway. The CCR4-NOT complex has been shown to mediate the codon usage effect on gene expression in yeast by affecting translation-dependent mRNA decay and mRNA solubility ^27,29,31^. Here we evaluated the role of this complex in human cells.

Depletion of CNOT4 from HEK293 cells indeed impaired the codon usage-dependent expression of our reporter genes, but it did not significantly affect the codon usage-dependent translation efficiency of mRNA reporters. Instead, we found that CNOT4 is highly enriched in nucleus and its depletion significantly affected the nascent and steady state nuclear levels and transcription of the reporter genes in a codon usage-dependent manner, indicating a role of CNOT4 in regulating transcription in a codon usage-dependent manner. Importantly, the effect of CNOT4 depletion on reporter mRNA level was maintained when translation was blocked. Although the CCR4-NOT complex functions in mRNA decay and translation in the cytoplasm, it was first identified by genetic selections for transcriptional regulators in yeast ^59,60^, and its nuclear functions in chromatin structure, transcription initiation and elongation, nuclear RNA quality control, and export have been previously described ^30^.

We previously showed that codon usage influences chromatin structure, and therefore transcription, in fungi, *Drosophila*, and human cells ^5,21,36^. Consistent with a role for CNOT4 in transcription, we found that depletion of CNOT4 affects mRNA levels genome-wide by preferentially upregulating mRNAs enriched with non-optimal codons. In addition, CNOT4 depletion also results in downregulation of nuclear mRNAs enriched for genes involved in chromatin regulation. The mechanism by which CNOT4 impacts gene transcription in a codon usage-dependent manner is not known. It is possible such an effect is caused by its impact on transcription of genes encoding for chromatin regulators. Consistent with this notion, we found that CNOT4 depletion result in significant downregulation of many chromatin regulators.

Furthermore, we previously showed that some chromatin regulators are involved in determining the codon usage effect on gene expression in *Neurospora* ^50^.

In addition to CNOT4, most of the top hits identified in our genome-wide screen are nuclear factors including multiple nuclear exosome components and subunits of its nuclear substrate adaptor, PAXT complex. Although the cytoplasmic exosome was expected to contribute to the codon usage effect by mediating translation-dependent mRNA decay, depletion of the nuclear exosome catalytic subunit DIS3, but not the cytoplasmic exosome catalytic subunit and its co-factors, significantly affected the codon usage-dependent expression of the reporter genes, indicating that the nuclear exosome has a larger impact on the regulation of genes with non-optimal codon usage profiles than does the cytoplasmic exosome.

ZFC3H1 and PABPN1 are PAXT complex subunits that associate with polyadenylated RNA substrates and target them to the nuclear exosome for decay ^46^. In human cells, depletion of either ZFC3H1 or PABPN1 significantly increased expression levels of the reporter genes with non-optimal codon usage. In addition, depletion of ZFC3H1 preferentially stabilized the mRNA with non-optimal codon usage with a concomitant rise in its cytoplasmic level. Furthermore, cytoplasmic and nuclear RNA sequencing of cells that were depleted with ZFC3H1 or PABPN1 revealed that PAXT is important for the global codon usage-dependent mRNA localization due to their roles in nuclear RNA quality control process. Interestingly, ZFC3H1 was previously shown to functionally compete with AlyREF, which is involved in recruitment of transcription/export (TREX) machinery to mRNA ^57^. TREX machinery has been shown to export primarily GC-rich mRNAs ^40^. It is possible that the specificity of TREX for GC-rich mRNAs is contributed by the exclusion of ZFC3H1 mRNA substrates from the TREX machinery.

Altogether, these results suggest that PAXT preferentially targets nascent mRNAs enriched with non-optimal codon usage for decay by the nuclear exosome. It was previously shown that codon usage and GC content of mRNA affect nuclear RNA export ^38–41,61^. Thus, different nuclear processes, including transcription, nuclear RNA quality control and mRNA export, are regulated by codon usage/nucleotide compositions, which preferentially promotes the expression and nuclear export of G/C-rich mRNAs with optimal codon usage and their subsequent translation in the cytoplasm. This conclusion is consistent with the recent hypotheses which proposed that high GC-content mRNAs, which are those with optimal codon usage, is a key mRNA feature that separates “wanted” transcripts from “unwanted” transcripts ^39,42,62,63^. The latter are mostly spurious or mis-spliced transcripts and RNAs transcribed from transposable elements or viruses. This hypothesis further proposes that in humans, a species with a small effective population size and long generation time, selection by translation is not prominent, instead much of the selection on human codon usage suppressed the production of “unwanted” transcripts through various nuclear effects to promote GC-rich transcripts and suppress AU/CG-rich ones for translation^62^. Although mRNAs of most genes with A/U-rich nucleotide composition or non-optimal codon usage are not “unwanted” mRNAs, their nucleotide composition-dependent regulation by transcription, nuclear RNA quality control and export pathways act to suppress their cytoplasmic levels to ensure their encoded proteins are not overexpressed to interfere with normal cellular functions.

Together, our results indicate that CNOT4, ZFC3H1 and PABPN1 differentially contribute to the nuclear codon usage effects on transcription, RNA quality control, and the preferential export of mRNAs with optimal codon usage. Several transcriptional regulators, such as TAFs and CCDC12, were also among the top hits identified in our CRISPR-Cas9 screen; however, it should be noted that the gene disruption-based CRISPR-Cas9 screen will likely miss many genes that encode factors involved in transcription and chromatin regulation due to their essential roles in normal cell growth and survival. Overall, these results demonstrate that the nuclear, translation-independent effects of codon usage have a large impact on gene expression levels in human cells.

## Acknowledgements

We thank Drs. Jaeil Han and Joshua Mendell for help in library screening; Vanessa Schmid in the McDermott Center Next Generation Sequencing Core for help in high throughput sequencing; Drs. Angela Mobley and Alyssa Guzman in the University of Texas Southwestern Flow Cytometry Core for assistance with cell sorting and Drs. Xueliang Lyu and Fangzhou Zhao for help in RNA sequencing analysis. We thank members of our laboratory for assistance and discussion. This work was supported by grants from National Institutes of Health (R35 GM118118) and the Welch Foundation (I-1560) to Y.L.

## Methods

### Cell culture, vector construction, transfection, and stable cell generation

HEK293T and HEK293 cells were cultured in Dulbecco’s Modified Eagle’s Medium (DMEM) supplemented with 10% fetal bovine serum (FBS) and 1% penicillin-streptomycin (Sigma, catalog #P4333). The cells were maintained in a humidified incubator at 37 °C with 5% CO_2_.

The dual reporter constructs, *CMV-mCherry^com^-CMV-GFP^rare^*and *CMV-mCherry^rare^- CMV-GFP^com^* ^45^, were a generous gift from Christopher M. Counter, Duke University. For the construction of *pCMV-wtGFP*, *pCMV-opt-GFP* and *pCMV-opt-mCherry* single reporter plasmids, PCR amplification was performed to generate fragments of *GFP^rare^*, *GFP^com^*, and *mCherry^com^,* using dual reporter constructs mentioned above as templates. Gel purified DNA fragments were digested with EcoRI and NotI and subcloned into pCMV-Tag-2B vector backbone at corresponding sites.

For transfection, polyethyleneimine transfection reagent, PEI (Polysciences, catalog #24765) was used according to the manufacturer’s instructions. Briefly, cells were seeded in wells of culture plates and allowed to adhere overnight. The transfection mixture was prepared by diluting the transfection reagents in Opti-MEM and adding DNA to the mixture. The mixtures were incubated at room temperature for 20 minutes and then added to the cells. After incubation, the transfection medium was replaced with fresh growth medium, and cells were allowed to recover before further analysis. The dual reporter constructs *CMV-mCherry^com^-CMV-GFP^rare^* and *CMV-mCherry^rare^-CMV-GFP^com^* were transfected into HEK293 cells followed by selection using G418 at 500 µg/mL concentration to generate cells that stably express wtGFP and opt-mCherry or opt-GFP and wt-mCherry. The stably transfected cells were sorted for dual fluorescence using FACS and diluted to generate clonal cell lines.

Co-transfection of single reporter plasmids was done as follows: *pCMV-wtGFP* or *pCMV-opt-GFP* was mixed with 1/10 amount of *pCMV-opt-mCherry* to normalize for transfection efficiency and diluted in Opti-MEM for transfection of HEK293T cells. The transfection mixture was prepared by diluting PEI in Opti-MEM and adding diluted DNA to the mixture. Transfections were performed as above and cells were harvested after 72 hours for western blot analyses.

### Generation of lentivirus and knockout cells

The lentiCRISPR_v2 vectors encoding Cas9, sgRNAs, and a puromycin-resistance gene and packaging plasmids were co-transfected into HEK293T cells at approximately 70% confluency in a 10-cm dish, and the medium was changed 24 hours later. After 48 hours of incubation, the cell culture supernatant was transferred to a 15-mL centrifuge tube and centrifuged at 3000×g for 10 minutes. The supernatant was then filtered through a 0.45-micron syringe filter and collected into a new sterile tube. The viral solution was further concentrated using the Lenti-X™ Concentrator (Takara, catalog #631231) according to the manufacturer’s instructions followed by snap freezing in liquid nitrogen and storage at -80 °C for long-term use.

Lentiviral sgRNA mediated knockout of genes was performed as described previously ^64^. HEK293 or HEK293T cells were split into media containing puromycin at a concentration of 1 µg/mL at 48 hours post-transduction. For experiments using *DIS3*-, *ZFC3H1*-, and *PABPN1*- knockout cells, selection in puromycin was done for 6-9 days before harvesting the cells for analysis. sgRNA sequences are provided in Table S1.

### siRNA and mRNA transfections

For siRNA-mediated knockdown, cells were transfected with 10 nM siRNA (Millipore Sigma) using Lipofectamine RNAiMAX reagent (Invitrogen) according to the manufacturer’s instructions. The details of siRNA sequences are provided in Table S1. For double knockdowns, 5 nM of each siRNA was used for a total mix of 10 nM. Cells were harvested at 72 hours post-transfection for all assays. The mRNA templates were prepared using *in vitro* transcription as previously described ^16^. mRNA transfections were performed using 0.5 µg of mRNA in 6-well plates using *Trans*IT-mRNA Transfection Kit (Mirus Bio) according to the manufacturer’s instructions, and cells were harvested after 8 hours.

### Immunofluorescence assay

HEK293 cells were transfected with 10 nM siRNA (Millipore Sigma) using Lipofectamine RNAiMAX reagent (Invitrogen) according to the manufacturer’s instruction. Cells were washed three times with phosphate-buffered saline (PBS) and fixed with 4% PFA for 20 min at 20°C at 72 hours post-transfection. Immunofluorescence assays were performed as previously described ^65^. The cells were analyzed using a Zeiss LSM 880 confocal microscope.

The sources of antibodies are listed in Table S1.

### Flow cytometry analysis

Cells stably transfected with reporter constructs were harvested and resuspended in PBS supplemented with 2% bovine serum albumin, 5 mM EDTA, and 0.05% sodium azide. Cells were filtered through a 35-micron filter and analyzed with a BD LSRFortessa cell analyzer. Data analysis was performed using FlowJo.

### Western blot analysis

The protein concentration of samples was determined by Bradford assay, and 10-80 µg of total, cytosolic, or nuclear protein extracts were separated by SDS-PAGE, transferred onto a PVDF membrane (Millipore), and detected using Pierce ECL western blotting substrate (Thermo Scientific, catalog #32106). The intensities of the bands were quantified using Fiji Image J software. The sources of antibodies are listed in Table S1.

### Reverse transcription and quantitative real-time PCR (RT-qPCR) assays

For analysis of gene expression and RNA subcellular localization, extracted RNA was treated with Turbo DNase (Ambion) and extracted using acidic phenol:chloroform (Ambion) prior to reverse transcription using a High-Capacity cDNA Reverse Transcription kit (Applied Biosystems) with random primers or HiScript III RT Supermix (Vazyme) according to the manufacturer’s protocol. Real-time PCR was performed as described previously ^5^. Primers used for RT-qPCR are listed in Table S1. For differential expression analysis, expression was normalized to *18S* rRNA or *GAPDH* mRNA. Fractionation quality was validated by primers targeting the nuclear *GAPDH-IN* pre-mRNA and the cytoplasmic *GAPDH* mRNA.

### Cellular fractionation

Cells were washed with PBS and trypsinised briefly. After stopping the reaction with DMEM, cells were collected by spinning at 400×g for 5 min. The cell pellets were resuspended in ice-cold PBS, and subcellular fractionation was performed as described previously ^66^.

Cytosolic and nuclear protein and RNA extractions were performed as previously described ^66^.

### Translation inhibition by cycloheximide

Cells were treated with siRNAs targeting *CNOT4* or with a control siRNA as described above. At 48 hours post-transfection, growth medium was replaced and cycloheximide (Sigma, catalog #C7698) dissolved in ethanol (10 mg/mL) was added to a final concentration of 100 µg/mL. Cells were harvested after 24 hours and RT-qPCR analyses were performed as above.

### Transcription inhibition by actinomycin D

Cells were treated with siRNAs targeting *ZFC3H1* or *CNOT4* or with a control siRNA in 10-cm plates as described above. At 72 hours post-transfection, growth medium was replaced and actinomycin D (Sigma, catalog #A9415) dissolved in dimethyl sulfoxide was added to a final concentration of 8 µg/mL. Cells were harvested at 0 hours (non-treated cells), after 1 hour, and after 2 hours of actinomycin D treatment. Cellular fractionations and RT-qPCR analyses were performed as above.

### Nuclear run-on assay

Cells were washed with cold PBS and collected by centrifugation at 500×g for 5 min at 4°C. Cells were lysed in 0.5 mL ice cold lysis buffer (10 mM Tris–HCl, pH 7.4, 10 mM NaCl, 3 mM MgCl2, 0.5% NP-40, and 10% Glycerol) supplemented with 200 U*/*mL Superase-in (Thermo Fisher) for 5 min on ice. Nuclei were isolated by centrifugation at 500×g for 5 min at 4°C and were resuspended in 50 µL freezing buffer (50 mM Tris–HCl, pH 7.4, 5 mM MgCl2, 40% glycerol and 0.1 mM EDTA) by pipetting. Transcription was performed by adding 50 µL transcriptional buffer (20 mM Tris–HCl, pH 8.0, 5 mM MgCl2, 300 mM KCl, 2 mM DTT, 500 µM ATP, 500 µM CTP, 500 µM GTP, 500 µM BrUTP (Sigma, B7166), 1% Sarkosyl and 200 U*/*mL Superase-in). After incubation at 30°C for 30 min (with shaking after every 5 min), 900 µL TRIzol was added to each reaction to stop transcription. RNA was isolated and resuspended in 100 µL IP buffer (50 mM Tris–HCl, pH 7.4, 150 mM NaCl, 0.05% NP-40, 1 mM EDTA, and 200 U*/*mL Superase-in), after removing an aliquot of 1/10 volume as input control. Anti-BrU antibody at 2 µg/IP (Santa Cruz Biotechnology, IIB5: sc-32323) and EZview Red Protein G beads (Sigma, E3403) were incubated with RNA for 3 hours at 4°C, followed by washing beads with IP buffer three times and isolation of RNA using TRIzol. Levels of newly transcribed RNA were measured by RT-qPCR.

### Genome-wide CRISPR-Cas9 screen

The Brunello lentiCRISPR_v2 library (Addgene, catalog #73179) ^67^ was used for genome-wide CRISPR-Cas9 screening as described previously ^68^. Two biological replicates were performed. Dual reporter-expressing HEK293 were spun down, resuspended in fresh media with 8 µg/mL polybrene (EMD Millipore), and mixed with the lentiviral library at a multiplicity of infection of ∼0.3. Beginning 48 hours after transduction, cells were selected in 1 µg/mL puromycin and grown for 10 more days before sorting. At least 2×10^7^ cells were transduced for each screen, corresponding to ∼300× or greater coverage. For cell sorting, cells were washed with PBS and resuspended in PBS supplemented with 3% FBS at 1.4 × 10^7^ cells/ml. The 0.5% of cells with the highest GFP to mCherry fluorescence ratio were sorted until 3 × 10^5^ cells were obtained per replicate. Cells were pelleted and frozen at -80 °C prior to genomic DNA extraction. At least 5 × 10^7^ unsorted cells per replicate were also collected and frozen. Genomic DNA from unsorted cells and sorted cells was extracted using DNeasy Blood & Tissue Kit (Qiagen, catalog #69504) according to manufacturer’s protocol.

For sequencing library preparation, two sequential rounds of PCR were performed using Herculase II Fusion DNA polymerase (Agilent) as described previously ^68^. The pooled PCR reaction mixture was used for the second PCR (9 to 11 cycles) with primers containing barcodes and adaptors for Illumina sequencing. Agencourt AMPure XP beads (Beckman Coulter Life Sciences) were used to purify amplicons. Library sequencing was performed on an Illumina NextSeq500 High Output sequencer to obtain 76-nucleotide single-end reads. Primer sequences are provided in Table S1. Approximately 400,000,000 sequencing reads were obtained for each replicate and MAGeCK was used to identify genes targeted by enriched sgRNAs in sorted populations as described ^69^.

### RNA-seq and data analysis

Knockdown efficiency was confirmed by FACS analysis. For total RNA-seq of si*CNOT4*- and siControl-treated cells, total RNA (from two biological replicates) was isolated using TRIzol reagent (Ambion). The RNA samples were then treated with Turbo DNase (Ambion) and extracted using acidic phenol:chloroform (Ambion). A poly-A RNA library was prepared using TruSeq Stranded mRNA Library Prep Kit (Illumina) following the manufacturer’s instructions, and sequencing was performed on the Illumina NextSeq 2000 sequencer to obtain 100-nucleotide single-end reads. Between 25,000,000 and 35,000,000 RNA-seq reads were mapped to the human genome (hg19 assembly). Fastq files were quality-checked using fastqc (v0.11.2). Reads from each sample were mapped to the reference genome using STAR (v2.5.3a)^70^. Read counts were generated using featureCounts ^71^, and differential expression analysis was performed using edgeR ^72^.

For the RNA-seq analysis of total, cytosolic, and nuclear RNA, cellular fractionation and RNA extractions were performed as above. Ribosomal RNA depletion and library preparation of total, cytosolic, and nuclear RNA were done using QIAseq FastSelect rRNA HMR (Qiagen) and UltraII Directional RNA Library Prep (NEB) kits following manufacturer’s protocols. The libraries were sequenced on Illumina NovaSeq X plus to obtain 60,000,000 (30,000,000 each strand) 150-nucleotide paired-end reads. FastQC (version v0.11.8) was used to check the quality of raw reads. Trimmomatic (version v0.38) was applied to remove adaptors and trim low-quality bases with default settings. RNA-seq reads were aligned to the human genome (hg38 assembly, GRCh38) with STAR Aligner version 2.7.1a, and Picard tools (version 2.20.4) were applied to mark duplicates. StringTie version 2.0.4 was used to assemble the RNA-seq alignments into potential transcripts, and featureCounts (version 1.6.0)/HTSeq was used to count mapped reads for genomic features such as genes, exons, promoter, gene bodies, genomic bins, and chromosomal locations. Differential expression was computed using DESeq2 (version 1.14.1), and the Gene Ontology analysis was done using ClusterProfiler package in R ^73^. The Venn Diagrams were drawn using Galaxy Version 0.1.1 ^74^.

For previously published total and nuclear RNA-seq data analyses, RNA-seq datasets were used from the study, GSE139151 (siNT total, cytosolic, and nuc samples in triplicates) ^40^. For GRO-seq data analyses, two different datasets from previously published studies were used ^44^ : Dataset 1: duplicate samples (DMSO_GRO-seq) of GSE136024. Dataset 2: duplicate samples (LP-1_DMSO) of GSE70449.

## Data availability

All high-throughput RNA sequencing data generated during this study have been deposited in Gene Expression Omnibus under accession numbers GSE261504 and GSE261505. All other data are available from the corresponding author upon reasonable request.

## Material availability

All materials and cell lines are available through University Texas Southwestern Medical Center from the corresponding author upon reasonable request.

## Contributions

R.G. and Y.L. conceptualized the study, analyzed and interpreted data and wrote the manuscript, which was revised and approved by all authors. R.G. and P.X. help designed and generated cell lines used in this study, R.G. J.D. and H.L. performed all cellular experiments, R.G. performed all bioinformatic analyses, and P.X. performed the immunofluorescent experiment.

---

**Figure S1.**
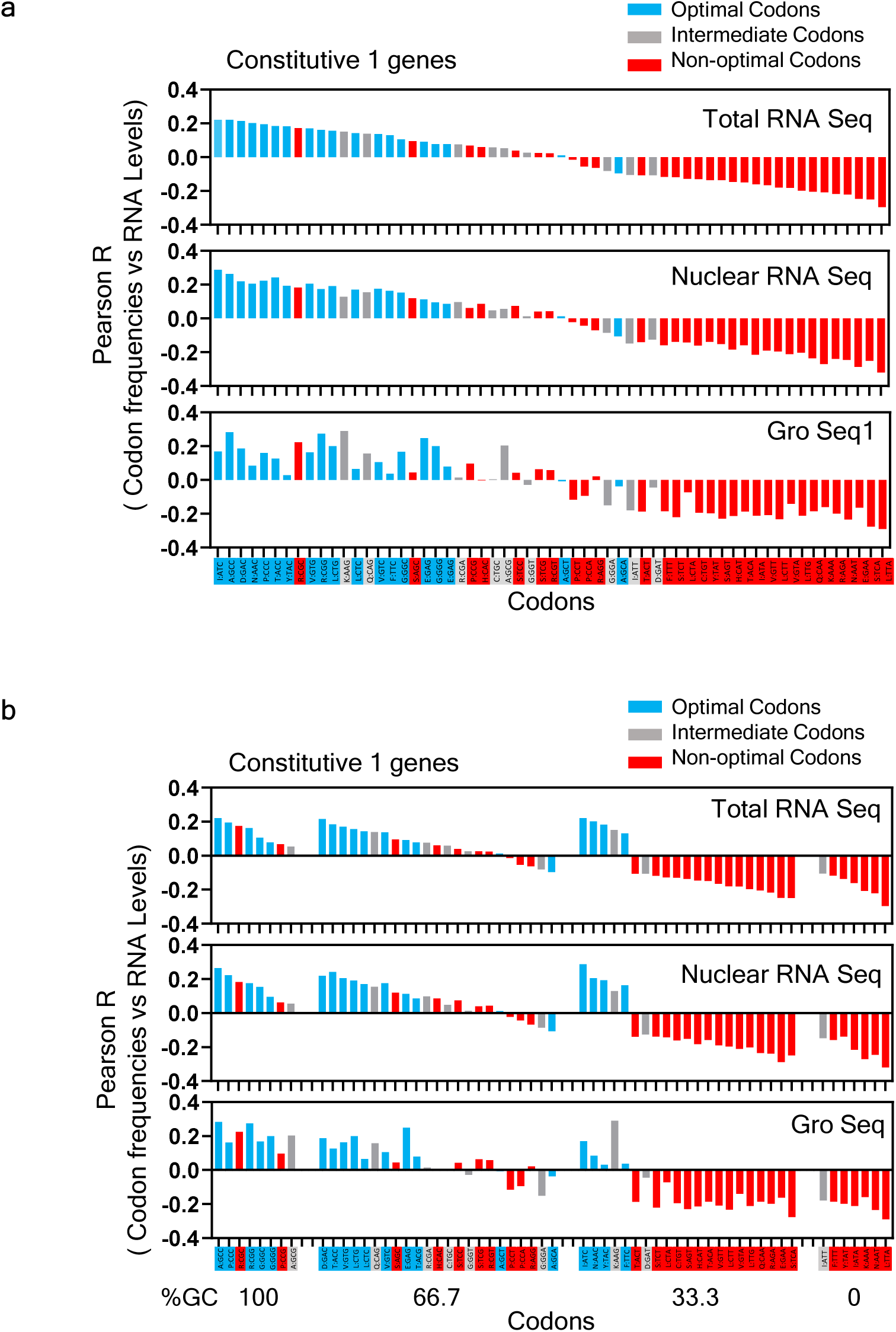
(a) Pearson correlation coefficients between occurrence of codon frequencies and RNA abundance for 59 synonymous codons in the total, and nuclear RNA samples from MCF1 cells and nascent RNA samples from HeLa cells (Gro Seq) calculated for constitutive1 genes are shown here. The correlation coefficients estimate the effect of codon optimality of each codon on transcript abundance in RNA samples. Optimal, intermediate, and non-optimal codons are indicated by blue, grey, and red bars, respectively. (b) The data in (Figure S1a) was replotted by grouping the codons based on their %GC contents and their correlation rankings as in (Figure S1a).

**Figure S2.**
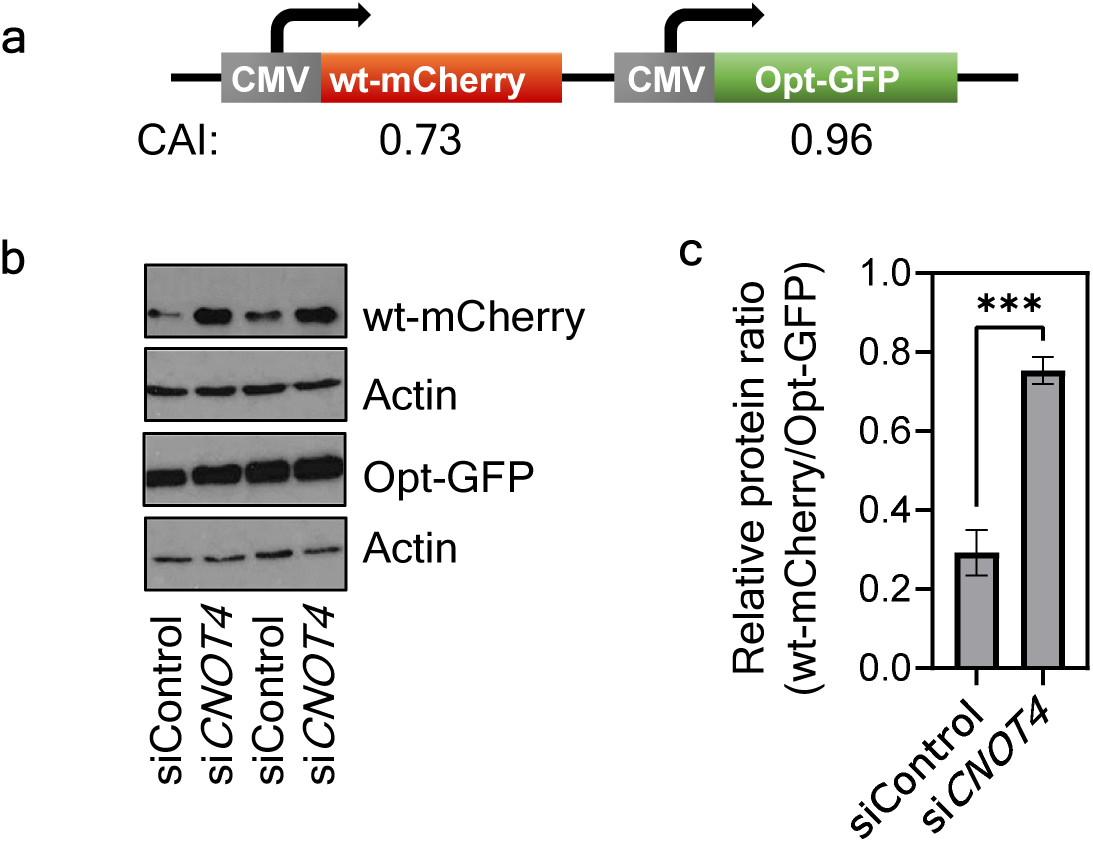
(a) Diagram showing the design of the CMV promoter driven wt-mCherry/opt-GFP dual codon usage reporter construct. (b) Western blot results showing opt-GFP and wt-mCherry protein levels in the wt-mCherry/opt-GFP stable cells treated with control siRNA or si*CNOT4* for 72 h. Actin protein levels are used as loading controls. (c) Densitometric analyses of Western blot results of opt-GFP and wt-mCherry protein levels as in (b) from three independent experiments. Data are presented as mean ±SEM, one-way ANOVA with Dunnett’s multiple comparisons test, ***P < 0.001.

**Figure S3.**
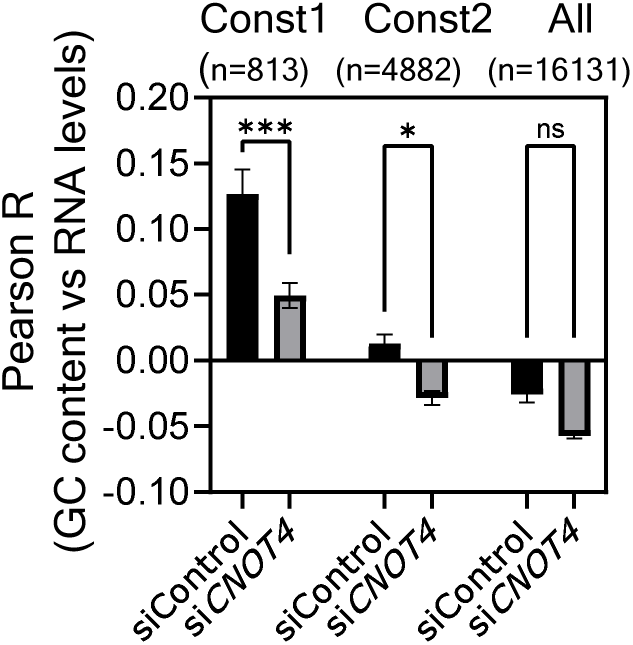
Plot showing Pearson correlation coefficients between GC content (nucleotide composition effect) and abundance of RNA in total RNA samples from HEK293 cells treated with either siControl or si*CNOT4* as indicated for 72 h followed by RNA sequencing, calculated for all genes or constitutive genes. Data is presented as mean ±SEM, n=2, one-way ANOVA with Sidak’s multiple comparisons test, ns - non significant; *P < 0.05; ***P < 0.001.

**Figure S4.**
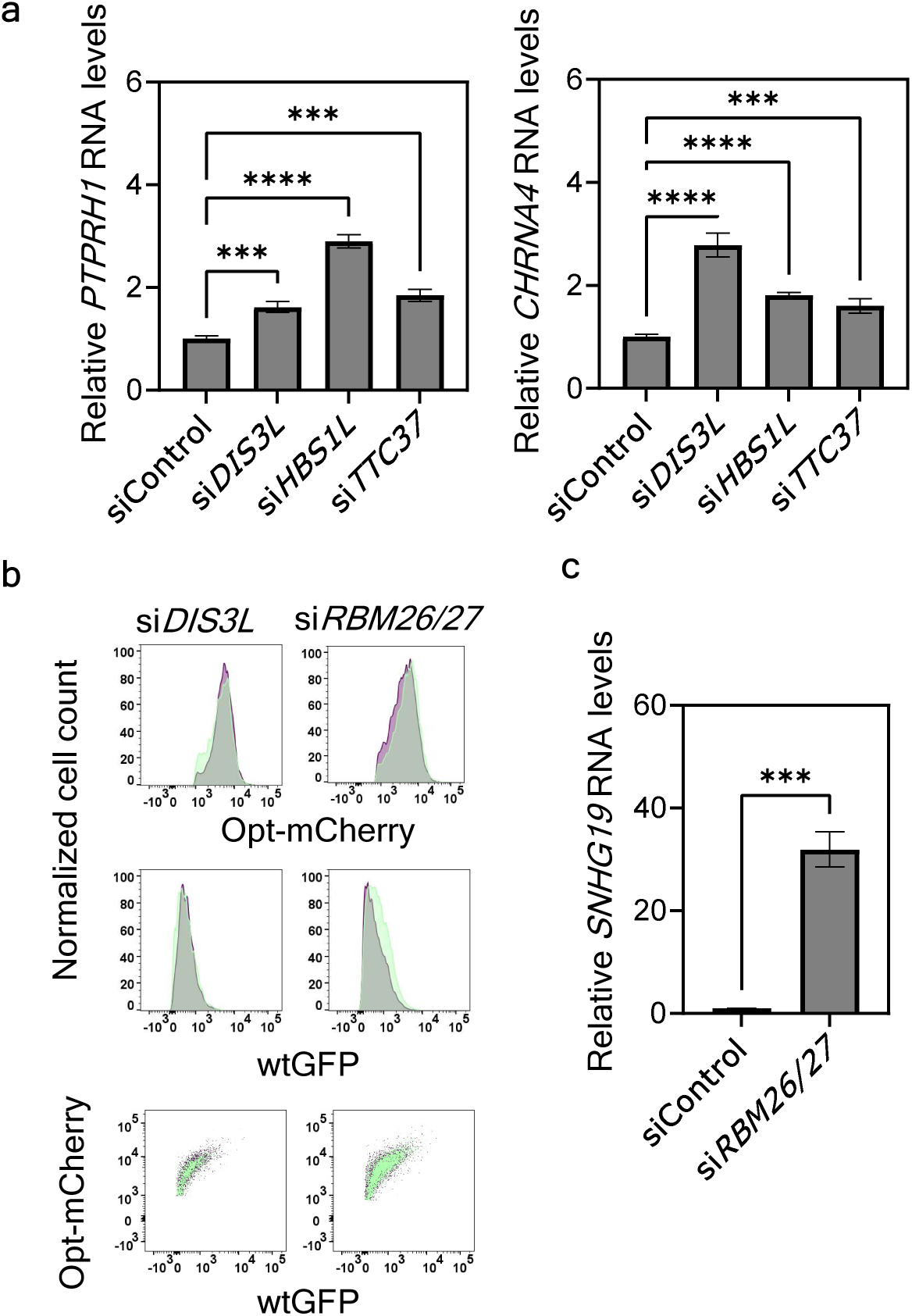
Involvement of nuclear but not cytosolic RNA exosome components in mediating codon usage effect on gene expression. (a) Quantification of the mRNA levels of *PTPRH1* (left) and *CHRNA4* (right) in total RNA isolated from wtGFP/opt-mCherry stable cells treated with indicated siRNA for 72 h as measured by RT-qPCR. Data are presented as mean ±SEM, n=3, one-way ANOVA with Dunnett’s multiple comparisons test, ***P < 0.001; ****P < 0.0001. (b) FACS analysis of wtGFP/opt-mCherry stable cells following siRNA mediated knockout of genes as indicated. Non targeting control siRNA treated cells are shown in green color while knockdown cells of DIS3L and RBM26/27 genes are shown in purple color in histograms and dot plots. (c) Quantification of *SNHG19* RNA levels in total RNA isolated from cells treated with control siRNA or si*RBM26/27* for 72 h. Data are presented as mean ±SEM, n=3, unpaired two-tailed Student’s t test, ***P < 0.001.

**Figure S5.**
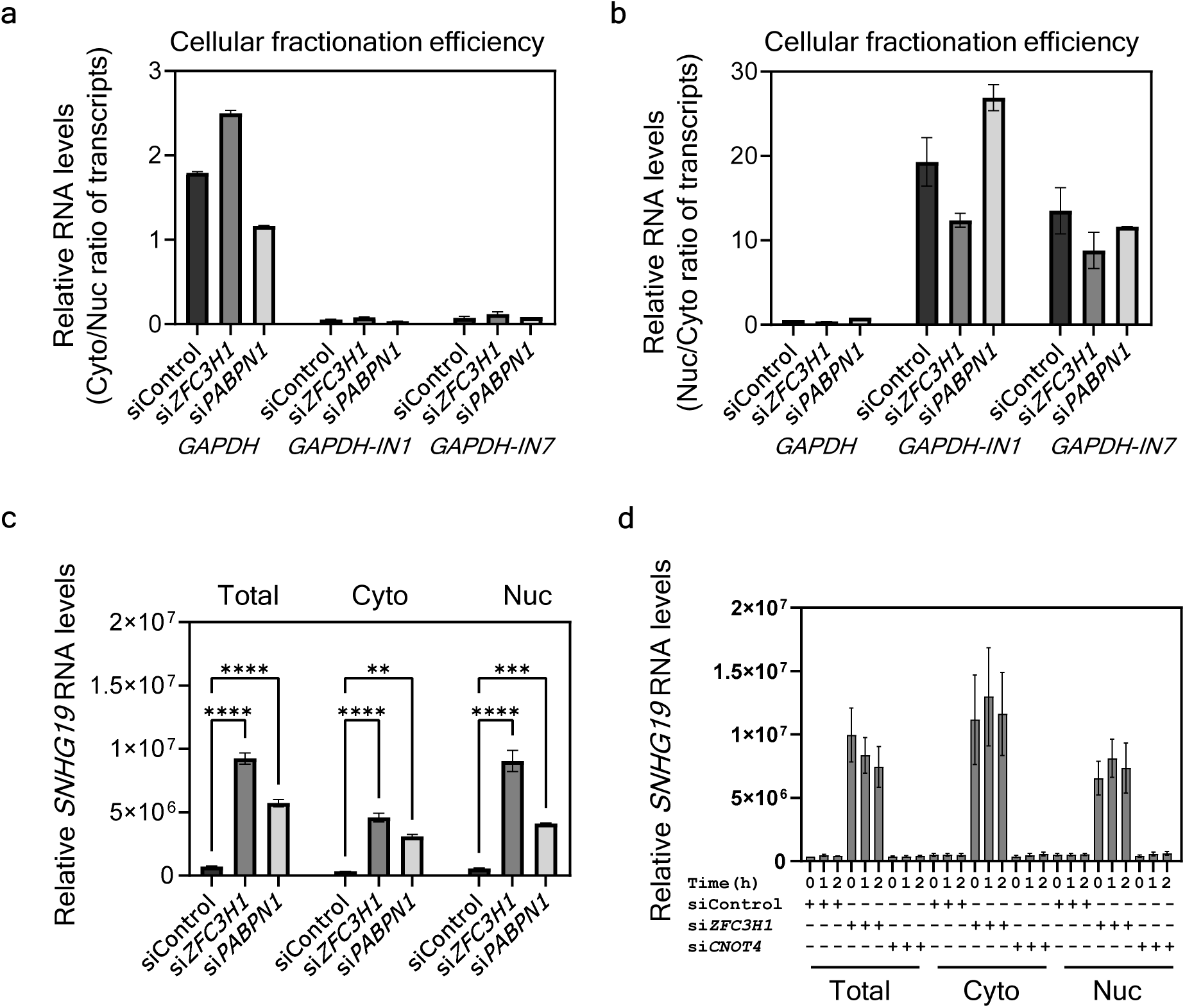
PAXT complex affects codon usage-dependent nuclear mRNA decay. (a-b) Quantification of the ratio of cytoplasmic to nuclear levels (a) or the ratio of nuclear to cytoplasmic levels (b) of *GAPDH* mRNA, *GAPDH-IN1* (intron1 containing) pre-mRNA and *GAPDH-IN7* (intron7 containing) pre-mRNA in RNA samples from Figure 6a as measured by RT-qPCR to determine cellular fractionation efficiency. (c) Quantification of *SNHG19* RNA levels in total, cytosolic and nuclear RNA samples from Figure 6a as measured by RT-qPCR. Data are presented as mean ±SEM, n=3, one-way ANOVA with Dunnett’s multiple comparisons test, **P < 0.01; ***P < 0.001; ****P < 0.0001. (d) Quantification of *SNHG19* RNA levels in total, cytosolic and nuclear RNA samples isolated from wtGFP/opt-mCherry stable cells subjected to indicated siRNA mediated knockdown for 72 h followed by actinomycin D treatment for different time intervals as in Figure 6c. RNA levels were measured by RT-qPCR.

**Figure S6.**
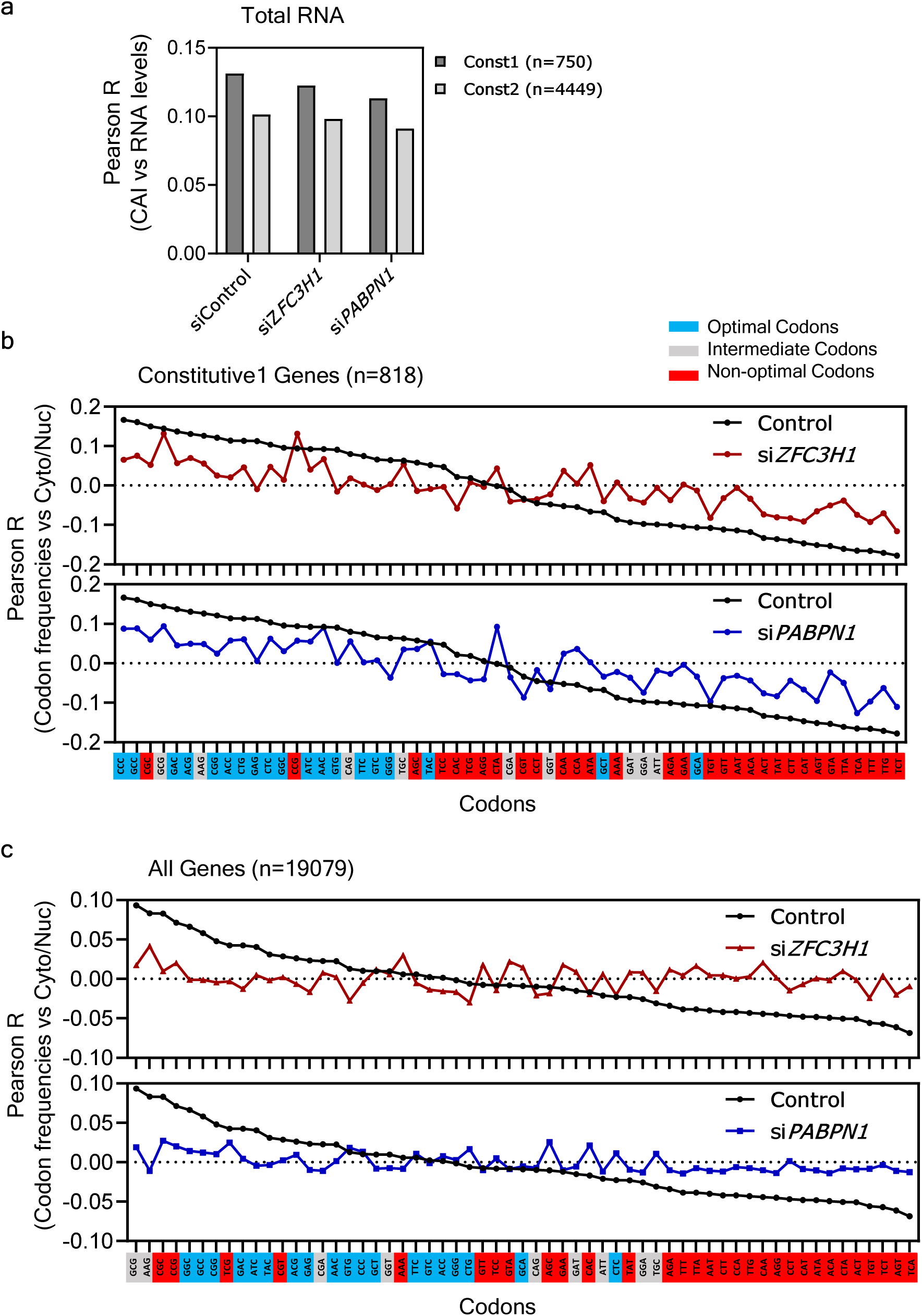
Genome-wide effects of PAXT complex components on codon usage mediated gene expression. (a) Plot showing Pearson correlation coefficients between CAI and total RNA levels in HEK293 cells treated with indicated siRNA for 72 h followed by RNA sequencing, calculated for constitutive1 and 2 genes (the numbers of genes in each group are shown in parenthesis). (b-c) Pearson correlation coefficients between occurrence of codon frequencies and cytoplasmic/nuclear ratio of mRNA for 59 synonymous codons in RNA samples from the control cells and the ZFC3H1 and PABPN1 knockdown cells for constitutive1 (b) or all genes (c) to estimate the effect of optimality of each codon on transcript abundance. Optimal, intermediate, and non-optimal codons are highlighted in blue, grey, and red colors, respectively. Number of genes (n) used in analyses in different groups are shown in parenthesis.

